# Splicing Retention and Enhancer Divergence Govern the Evolutionary Fate of Ohnologues Following Whole-Genome Duplication in Rainbow Trout

**DOI:** 10.1101/2025.08.06.669003

**Authors:** Ali Ali, Rafet Al-Tobasei, Huaijun Zhou, Mohamed Salem

**Affiliations:** Department of Animal and Avian Sciences, University of Maryland, College Park, College Park, MD, USA; Computational Science Program, Middle Tennessee State University, Murfreesboro, TN, 37132, USA; Department of Animal Science, University of California, Davis, Davis, CA 95616, USA

## Abstract

Whole-genome duplication (WGD) produces gene duplicates—known as ohnologues—that expand evolutionary potential while posing challenges for regulatory coordination and functional balance. While divergence in gene expression has been widely studied post-WGD, the long-term evolutionary dynamics of alternative splicing (AS) and its regulatory basis remain unresolved. Here, we investigate the evolution of AS in rainbow trout (Oncorhynchus mykiss), a species that underwent a salmonid-specific WGD ∼100 million years ago. Using a high-quality genome assembly, transcriptome data across six tissues, and ChIP-seq profiling of histone modifications, we classify ohnologue pairs based on their expression divergence, splicing complexity, and epigenetic signatures. We find that most ohnologues are retained through conservation, with a gradual reduction in AS diversity over time. Contrary to earlier models, a substantial fraction of ohnologues follows an independent splicing model, maintaining splicing complexity similar to their unduplicated ancestors. Strikingly, we show that enhancer-associated histone marks, particularly H3K27ac, diverge significantly between neofunctionalized and independently splicing gene pairs, implicating enhancer rewiring as a key driver of regulatory and functional divergence. These findings reveal that AS evolution after WGD is shaped by both selective pressures and epigenetic modulation, challenging assumptions of rapid splicing loss and highlighting the independent model as a dominant long-term fate. Our results provide a unified framework for understanding how splicing and regulatory landscapes evolve following genome duplication, with broad implications for vertebrate genome evolution and functional innovation.

## INTRODUCTION

Whole genome duplication (WGD) increases functional redundancy and genomic complexity in evolving eukaryotes[1, 2]. The divergence of gene expression and alternative splicing (AS) forms following WGD has been the subject of a significant and ongoing debate. For instance, the relative importance of sub– and neo-functionalization in the genome evolution following WGD events has been controversial. Neofunctionalization was previously reported to be encountered less frequently than subfunctionalization in fish[3]. In agreement, the sum of expression levels of duplicated teleost genes supported the subfunctionalization model[4]. Conversely, a recent study has found neofunctionalization to be far more common than subfunctionalization in Atlantic salmon[5]. Sandve et al. reanalyzed data from the two contradictory studies[4, 5] and found that the discrepancy resulted from the analytical approach[6]. By ranking the individual genes within the duplicate pairs, the study concluded that neofunctionalization is likely the prevailing fate for ohnologues from the third teleost-specific (Ts3R) and the salmonid-specific fourth vertebrate (Ss4R) WGD events[6]. The study by Sandve et al. evaluated the global pattern of gene expression evolution; however, the analyses did not consider that selection acts differently on individual genes[7]. Additionally, Braasch et al. pointed out that pooling together ohnologues with the most correlated expression patterns automatically results in them being more correlated with the unduplicated sister taxon than the less conserved ohnologues[7]. Thus, further studies of additional fish genome duplication events and new bioinformatics tools were recommended to evaluate the prevailing fate for ohnologues on a gene-by-gene basis while comparing gene pair expressions to singletons in pre-duplication ancestors[6, 7].

Further, the evolutionary relationship between AS and gene duplication has long been debated. Initially, it was established that an inverse correlation exists between the size of the gene family and the fraction of genes with AS forms[8, 9]. A more recent study revealed that the relationship between gene family size and the proportion of genes with AS forms is not inverse but exhibits a parabolic pattern[10]. The trend can be more precisely elucidated by stating that lower levels of splicing are observed in single-copy genes and genes with family sizes above 10, as opposed to those between 2 and 10[10]. The evidence further demonstrated that duplications occurring in the distant past had higher proportions of genes with AS forms than more recent ones[8]. A consistent hypothesis suggested that splicing forms are lost almost immediately after a gene duplication event[9]. Thus, it was proposed that this finding of a reduction in the number of AS forms at the genome level aligns more closely with the predictions of the function-sharing model rather than the independent model[9]. Additionally, a previous study[11] suggested a combined scenario of function-sharing and accelerated models for the evolution of AS in teleost’s ohnologues while ruling out the independent model. The fact that splicing levels change following duplication suggests a potential interdependence between the two evolutionary mechanisms. Despite multiple salmonid genome assemblies, the evolutionary trajectory of salmonids AS and its relationship with gene duplication remains not fully explored.

In this study, we used our recent well-assembled rainbow trout (*Oncorhynchus mykiss*) reference genome (Swanson doubled haploid genetic line) to investigate the early evolutionary fate of duplicated gene copies and AS forms following WGD. We discovered higher levels of conservation than previously thought in a closely related species, the Atlantic salmon[5]. We also found that neofunctionalized genes are evolving rapidly at synonymous sites, which may reflect relaxed selective constraints. Conserved genes hold more AS forms than neofunctionalized and specialized genes, partly influenced by their ancestral orthologs’ splicing level. Furthermore, we concluded that ohnologues gradually lose AS forms with evolutionary time, and we introduced the independent model as the primary long-term evolutionary model of AS following WGD events. This finding challenges the conclusions drawn by a previous study[11], which had utterly dismissed the independent model’s role in the evolutionary process of teleost’s AS. Finally, the distinctive epigenetic profiles observed in neofunctionalized and independent gene pairs reflect their evolving regulatory mechanisms, supporting the idea that epigenetic divergence plays a key role in regulating splicing and driving functional innovation.

## RESULTS

### Evolutionary processes driving preservation of gene duplicates

The ancestral genome of all teleost fish underwent three rounds of whole genome duplication (WGD). However, salmonids experienced an additional 4th WGD around 80-100 million years ago. Therefore, the rainbow trout genome provides a special opportunity to study which genes are retained or lost, gain or lose splice variants, and the evolutionary mechanisms that maintain duplicate genes and alternative splicing forms.

The mechanism of duplicate gene retention has been controversial. Rapid expression divergence has been widely acknowledged as a major factor in the preservation of duplicated genes[12–15], along with a significant debate regarding the relative contribution of both sub– and neofunctionalization[4–7]. Therefore, we set out to investigate the evolution of gene expression in rainbow trout across six tissues (brain, liver, muscle, intestine, kidney, and spleen) (Figure 1A-E).

**Figure 1.**
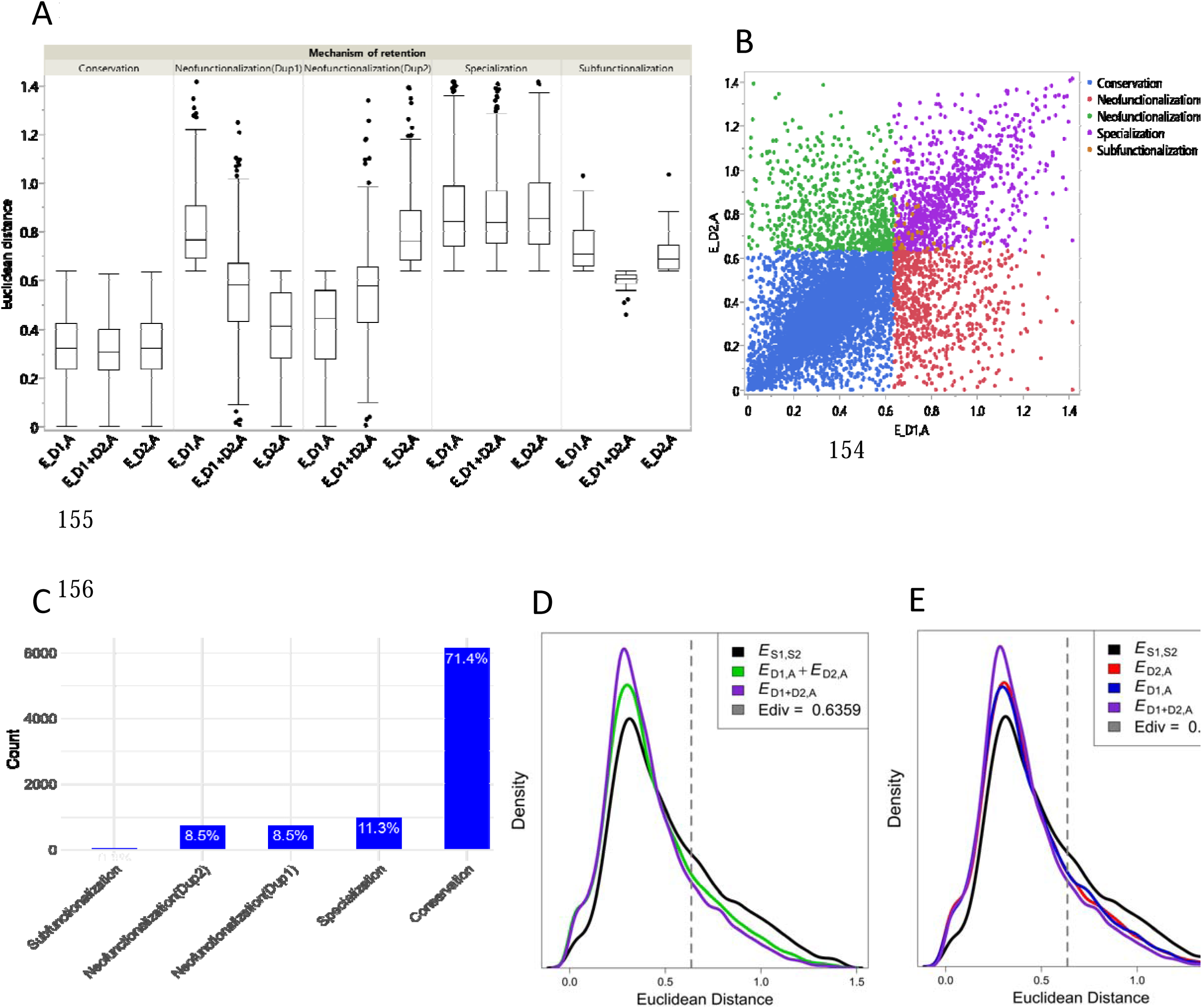
(A) Classification of evolutionary processes preserving duplicate gene copies. Gene pairs with *E*_D1,A_ ≤ 0.6359 and *E*_D2,A_ ≤ 0. 6359, were classified as conserved genes. Gene pairs with *E*_D1,A_ > 0. 6359 and *E*_D2,A_ ≤ 0. 6359 or *E*_D1,A_ ≤ 0. 6359 and *E*_D2,A_ > 0. 6359 were classified as neo-functionalized. Under sub-functionalization, *E*_D1,A_ > 0. 6359, *E*_D2,A_ > 0. 6359, and *E*_D1+D2,A_ ≤ 0. 6359 were expected. Specialized genes had *E*_D1,A_ > 0. 6359, *E*_D2,A_ > 0. 6359, and *E*_D1+D2,A_ > 0. 6359. (B) The plot shows Euclidian distances between expression profiles of D1 and ancestral copies (*E*_D1_,_A_) on the x-axis and Euclidian distances between expression profiles of D2 and ancestral copies (*E*_D2_,_A_) on the y-axis. Blue, red, green, purple, and brown dots represent conserved, neo-functionalized (D1), neo-functionalized (D2), specialized, and subfunctionalized genes, respectively. (C) Count and percentage of duplicate gene copies retained through various mechanisms. (D & E) Distributions of all computed Euclidian distances and the position of E_div_. *E*_D1,A_ + *E*_D2,A_ is a single distribution for *E*_D1,A_ and *E*_D2,A_. The plots suggest that most of the Ss4R ohnologues are mainly retained by conservation.

Accurate quantification of duplicate gene expression can be challenging when sequencing reads map to both gene copies or if the latter have differential homology with other locations in the genome[16]. Therefore, we estimated gene expression levels using reads uniquely mappable to each paralogous position. The same approach was previously used for measuring expression ratios of duplicate genes in mammals[16]. Random order was given to the duplicate gene copies (i.e., labels duplicate 1 “D1 or Dup1” and duplicate 2 “D2 or Dup2” were randomly designated to the two genes). We identified 9,701 ohnologue pairs corresponding to 7,422 singletons in the Northern pike. Among them, 8,596 triplet gene copies were found, with each copy being expressed in at least one tissue.

To distinguish the evolutionary processes driving the preservation of gene duplicates following WGD (Figure 1A-E), we adopted the phylogenetic approach developed by Assis and Bachtrog[17, 18] (See Methods Section). We quantified the divergence of gene expression profiles among the rainbow trout’s duplicate pairs and ancestral genes in the Northern pike. In this study, we used the Euclidian distances method, not the Pearson’s correlation coefficient, because of its robustness to measurement error and low estimates of expression divergence for genes with conserved uniform patterns of expression[17, 19]. Raw/absolute expression levels were converted to relative expression values before calculating the Euclidean distance[17, 19]. We computed Euclidian distances between the expression profiles of D1 and ancestral copies (*E*_D1,A_), D2 and ancestral copies (*E*_D2,A_), and sum D1-D2 expression profiles and corresponding ancestral copies (*E*_D1+D2,A_). Also, we computed Euclidian distances between expression profiles of all singletons in Swanson rainbow trout and Northern pike (*E*_S1,S2_) and used it as a baseline divergence level for genes[17, 18]. Duplicate genes were classified using previously defined rules[17] (Figure 1A). We identified 71.4% cases of conservation, 17.0% cases of neofunctionalization, 11.3% cases of specialization, and 0.3% cases of subfunctionalization (Table S1). Our analysis showed that most duplicates are maintained through conservation (Figure 1B-E). Neofunctionalization and specialization were observed in smaller percentages of the WGD genes, and subfunctionalization was rare.

Ranking the individual genes within the duplicate pairs[6] led to a discrepancy in the main conclusion of two previous studies[4, 5] investigating the mechanism of gene duplicate preservation. To test whether the ranking would affect our conclusion, we ranked the gene pairs as “most conserved” (i.e., the gene copy with the highest expression correlation with the ortholog in the outgroup; D1) and “most diverged” (i.e., the gene copy with the lowest expression correlation with the ortholog in the outgroup; D2) as in[6]. Interestingly, our approach was robust to the ranking process and yielded the same results from random ranking (Table S2).

Duplicate genes may show tissue-specific, time-point-specific, or simultaneous divergent expression[20]. Therefore, we sought to replicate our conclusion using RNA-seq data from fish used to assemble the rainbow trout[21] (BioProject PRJEB4450) and Atlantic salmon[5] (BioProject PRJNA72713) genomes. Consistent with the results above, more than 70% of duplicate genes were preserved by conservation (Figure S1 and Tables S3 & S4). To further validate our results and eliminate potential bias caused by the developmental, physiological, and immunological conditions of the studied fish, we reanalyzed the data using house-keeping genes that displayed TPM values ≥ 1 in each tissue of both the Northern pike and rainbow trout [22] (BioProject PRJNA389609). Consistent with the whole dataset, a similar pattern was also achieved when only housekeeping genes were used (69% conservation, 16.6% neofunctionalization, 14.0% specialization, and 0.4% subfunctionalization) (Table S5). Taken together, we have discovered higher levels of gene conservation than previously thought in salmonids based on a previous Atlantic salmon study [5] (42%). Additionally, our findings demonstrate that the relative significance of sub– and neo-functionalization aligns with a recent study conducted on Atlantic salmon[5].

#### Long-term preservation of duplicate gene pairs

We hypothesized that non-salmonid teleost fish that have undergone only Ts3R event would offer a long-term picture of gene evolution. To test this hypothesis, RNA-Seq data from pike, zebrafish, and medaka were analyzed to quantify the divergence of gene expression profiles among duplicate pairs and corresponding ancestral genes in gar (outgroup). Due to the limited signature of the Ts3R event, we identified a smaller number of ohnologues corresponding to singleton orthologs in the spotted gar (1863, 740, and 858 triplets in Northern pike, zebrafish, and medaka, respectively) compared to the Ss4R ohnologues. Particularly in medaka, we have noticed a decrease in the proportion of conserved genes, while there has been an increase in the proportion of neofunctionalized genes when compared to salmonid fish (Figure 2A-C and Table S6). In contrast, a comparable proportion of specialized genes was observed in both the Ss4R and non-salmonid ohnologues. Notably, the Ss4R neofunctionalized genes revealed a significantly higher average of dS (number of substitutions at synonymous sites) than other duplicate gene types (Figure 2D). Altogether, our findings suggest that neofunctionalized genes are evolving more rapidly at synonymous sites, which may reflect relaxed selective constraints or increased mutation rates.

**Figure 2:**
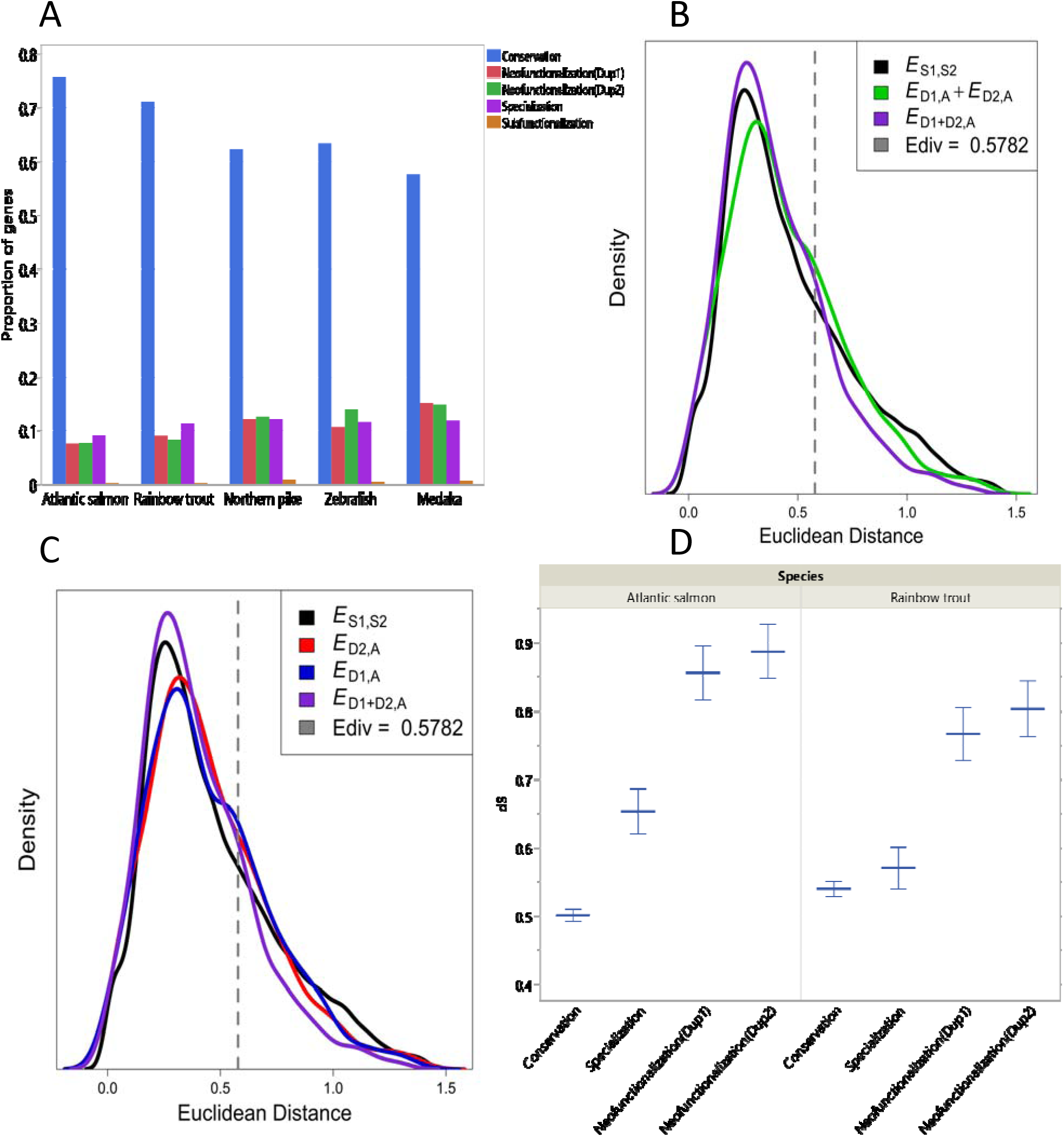
Evolutionary processes retaining duplicate gene copies after the Ss4R and Ts3R events. (A) The proportion of conserved gene pairs decreased in non-salmonid fish while the proportion of neo-functionalized genes increased. (B & C) In medaka, a rightward shift in the distribution of *E*_D2,A,_ *E*_D2,A,_ and their single distribution *E*_D1,A_ + *E*_D2,A_ suggests that a significant portion of duplicate gene pairs in medaka diverged in expression from their singleton orthologs. *E*_D1,A_ represents Euclidian distances between the expression profiles of D1 and ancestral copies, whereas *E*_D2,A_ represents Euclidian distances between the expression profiles of D2 and ancestral copies. *E*_D1,A_ + *E*_D2,A_ is a single distribution for *E*_D1,A_ and *E*_D2,A_. (D) In rainbow trout and Atlantic salmon, the Ss4R ohnologues likely tend to neofunctionalize as the evolutionary time (dS) progresses.

### Evolution of alternative splicing following WGD

#### Ohnologues do not lose AS forms immediately following WGD

In our analysis of 38,903 chromosome-anchored trout genes, we discovered that 31,674 of them (81.4%) are alternatively spliced, meaning they may have at least one alternatively spliced form (i.e., two forms of mRNA). There are typically 7.3 different AS isoforms per trout gene. Notably, 7,856 genes (20.2%) have >10 AS forms. We noticed that the number of genes decreases as the number of AS forms/gene increases (Figure 3A).

**Figure 3:**
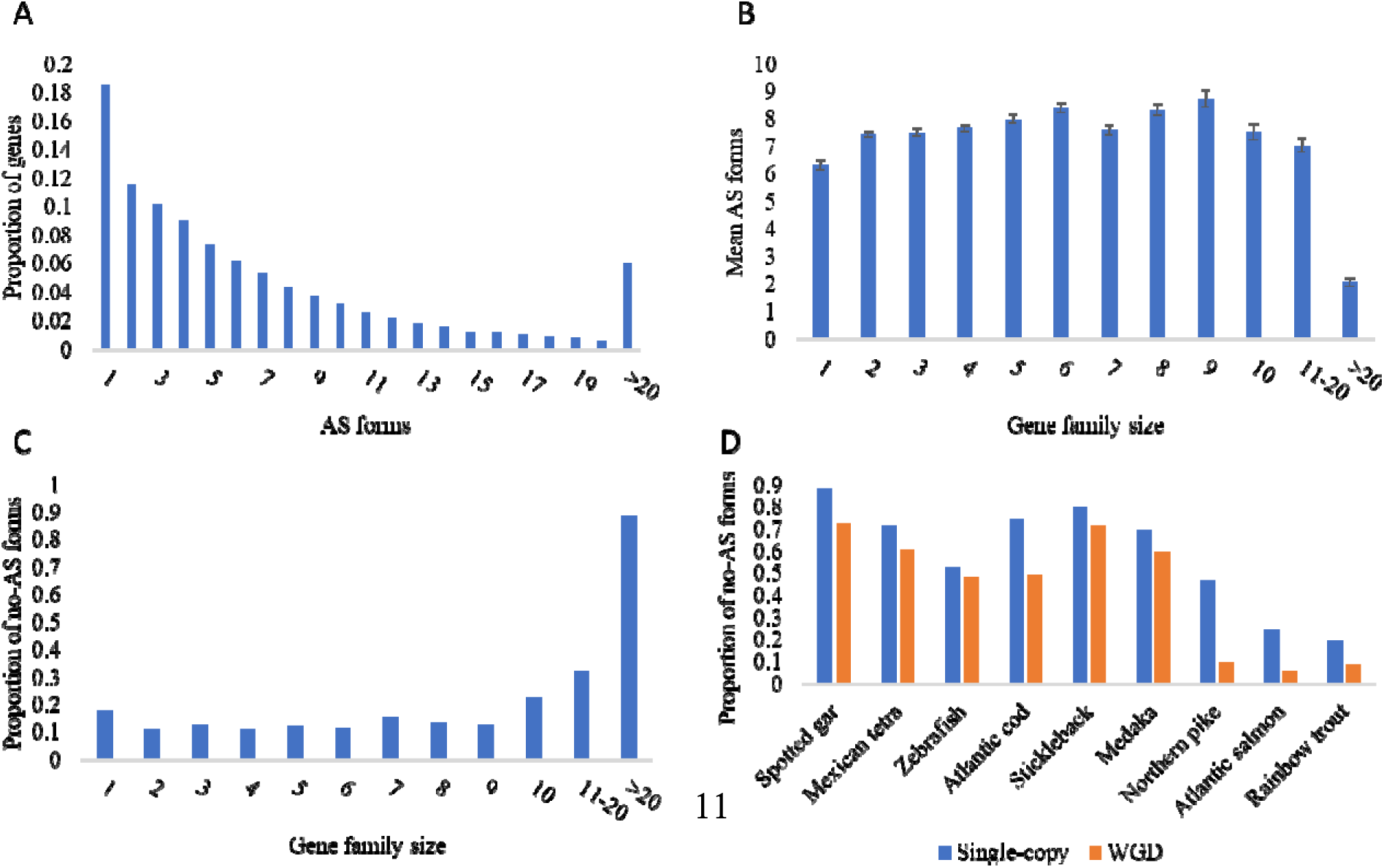
Alternative splicing and gene duplication are not inversely correlated. (A) Distribution of AS forms in the rainbow trout genes. Each bar shows the proportion of trout genes that have a specific number of AS forms. (B) Larger gene families have more AS forms than singletons. (C) The plot shows the fraction of genes not alternatively spliced on the y-axis and the trout gene family size on the x-axis. (D) WGD duplicates have fewer no-AS forms than single-copy genes in salmonid and non-salmonid teleost fish. The proportion of no-AS forms in WGD duplicates ranged from 0.73 in spotted gar, 0.06 in Atlantic salmon, and 0.09 in rainbow trout.

The relationship between gene duplication and AS has been the subject of a substantial debate for approximately two decades. Evidence demonstrated that duplications occurring in the distant past had higher proportions of genes with AS forms than more recent ones[8]. A consistent hypothesis suggested that splicing forms are lost almost immediately after a gene duplication event[9]. To test the hypothesis proposed by Su et al in rainbow trout, we performed all-by-all BLASTp to find the most similar protein sequences, which were clustered into 12,092 gene families. Gene families were categorized into 12 gene family sizes (Figure 3B & C). For instance, gene family size 1 has all single-copy genes, whereas gene family size >20 has all families with more than 20 genes. A parabola-like curve was observed between the mean AS forms and gene family size (Figure 3B). The mean number of AS forms in duplicated gene families, except gene family size >20, was higher than in the single-copy gene family. Single-copy genes have significantly fewer AS forms (82.3%) than genes of family sizes two to nine (87.8%) (χ2 test, P = 0). In particular, gene families containing exactly two members exhibited a higher number of AS events compared to single-copy genes (Figure 3B & C and Figures S2-S9). While there is a higher proportion of alternatively spliced genes in larger gene families, a remarkable increase in the proportion of the genes with no alternative splicing (no-AS) compared to single-copy genes was observed only in gene family sizes ≥ 10 (49.6%) (χ2 test, P = 2.68E-172) (Figure 3C). Overall, our results revealed that the relationship between gene family size and the proportion of genes with AS forms is not inverse and that gains of AS forms accompany gene duplication.

We also noticed that genes in colinear blocks (WGD duplicates) tended to have fewer no-AS form than single-copy genes across evolutionary-related fish species (Figure 3D). The mean number of AS forms per single-copy genes is 6.33 ± 0.16, which is significantly lower than that of WGD duplicates (8.28 ± 0.06) (Wilcoxon rank sum test, *P* < 2.2e-16). The mean number of AS forms highly correlated (R = 0.87) with the proportion of WGD duplicates across all genomes investigated in this study (Wilcoxon signed rank test, *P* = 0.002). According to Su et al.[9], we roughly estimated the acquisition of AS form due to gene duplications by multiplying the number of duplicate genes by the difference in mean between single-copy and duplicated genes. We found gains of ∼47,514 AS forms (24,366 x 8.28 – 6.33 = 47,514) due to WGD in the rainbow trout genome.

Further, 10,522 Ss4R ohnologue pairs (20,915 genes) were used to investigate the relationship between the evolutionary time (dS) and the proportion of no-AS forms following WGD. Notably, we detected a positive correlation between the dS bins and the no-AS forms (R = 0.87, *P* = 0.005) (Figure 4A), contradicting the notion that AS forms are lost immediately after duplication[9]. Conversely, we detected a negative correlation between the evolutionary time of dispersed duplicate genes (19,036 gene pairs) and no-AS forms (R = –0.74, *P* = 0.03) (Figure 4B), supporting the notion that duplicated genes acquired more AS forms with the time. Thus, our results demonstrate that ohnologues lose splice variants with evolutionary time under relaxed purifying selection pressure, in contrast to the dispersed duplicates.

**Figure 4:**
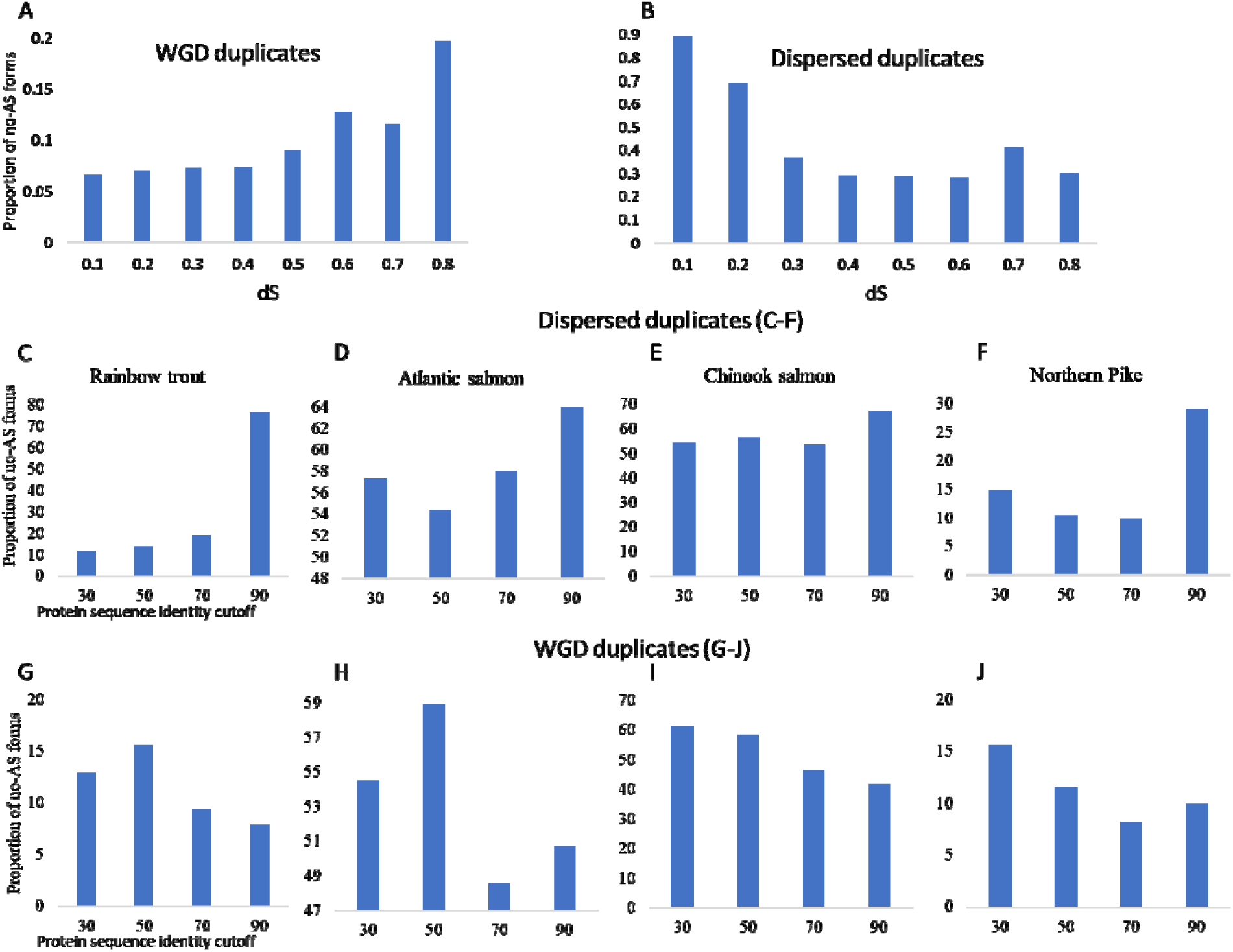
(A) A positive correlation between the protein sequence identity and proportion of no-AS forms supports the notion that AS forms are lost as the WGD duplicates evolve with time. (B) A negative correlation between the protein sequence identity and proportion of no-AS forms indicates the acquisition of AS forms as the dispersed duplicates age. Recent dispersed duplicates are less likely to be alternatively spliced (C-F), whereas WGD duplicates tend to be less alternatively spliced as they evolutionary diverge (G-J). Each bar represents the fraction of genes not subjected to splicing in four evolutionary-related fish species. Group 30 consisted of genes showing a sequence identity ranging from 30% to 50% (30 ≤ *I* < 50) at an E-value less than 1E-5. Similarly, the groups 50, 70, and 90 had sequence identity within the ranges of 50 ≤ *I* < 70, 70 ≤ *I* < 90, and *I* ≥ 90, respectively. The titles of the x– and y-axes in the leftmost panel apply to the panels on the right as well.

The above analysis used trout AS isoforms (Figure 4A & B). Thus, we sought to further investigate in other fish whether the Ss4R ohnologues and dispersed duplicates also tend to have reciprocal patterns of AS form acquisition as they evolve. We used the method Su et al. (2006)[9] developed to compare the evolutionary patterns of AS in rainbow trout and other fish species (Figure 4C-J). In this approach, we used the amino acid identity percentage (*I*), at an E-value less than 1E-5, from the self-genome BLASTp output to group the duplicate genes into four bins (30 ≤ *I* < 50, 50 ≤ *I* < 70, 70 ≤ *I* < 90, and *I* ≥ 90). We then calculated the proportion of genes with no-AS forms in each bin. The amino acid identity (*I*) was used as a proxy for the time of divergence of duplicate gene copies. Thus, it is expected that as *I* increase, the proportion of genes with no-AS forms will decrease if the loss of AS isoforms in WGD duplicates does not happen shortly after the gene duplication. Also, it would be expected that as *I* increase, the proportion of genes with no-AS forms will increase if the loss of AS isoforms in dispersed duplicates happens shortly after the gene duplication. Figure 4 shows this was the case in rainbow trout, Atlantic salmon, chinook salmon, and Northern pike. A consistent pattern was also observed in coho salmon (Figure S10). Overall, ohnologues have stronger selective pressure to maintain functional redundancy, potentially leading to the retention of AS forms. As selective pressure decreases with greater divergence, AS forms may be lost. Our analysis suggests that the loss of AS forms is contingent upon the degree of divergence between duplicate genes, selective pressure and functional redundancy, and type of duplication (WGD vs. dispersed duplicates).

Furthermore, we found that conserved genes hold more AS forms than neofunctionalized and specialized genes, suggesting a link between gene conservation and splicing complexity. Conserved gene pairs, which exhibit higher ancestral splicing, likely help maintain gene dosage balance (Figure 5A). In contrast, neofunctionalized and specialized genes show a reduction in splicing levels, reflecting evolutionary shifts toward new and distinct functions (Figure 5A). The pairwise comparison revealed a significant difference between the mean ancestral splicing of all possible pairs of conserved, neofunctionalized, and specialized genes. For instance, conserved and specialized genes’ mean ancestral splicing levels were 9.8 and 7.8, respectively (Wilcoxon test, *P* = 7.8E-17). Additionally, we have observed a highly significant difference in the expression levels of the conserved duplicates compared to the neofunctionalized (Wilcoxon test, P < 2.5E-61) and specialized genes (Wilcoxon test, P < 2.3E-42) (Figure 5B). The significant differences in splicing forms and expression levels between conserved, neofunctionalized, and specialized genes further underscore the role of AS in maintaining functional balance in conserved genes, while allowing evolutionary innovation in neofunctionalized and specialized duplicates.

**Figure 5:**
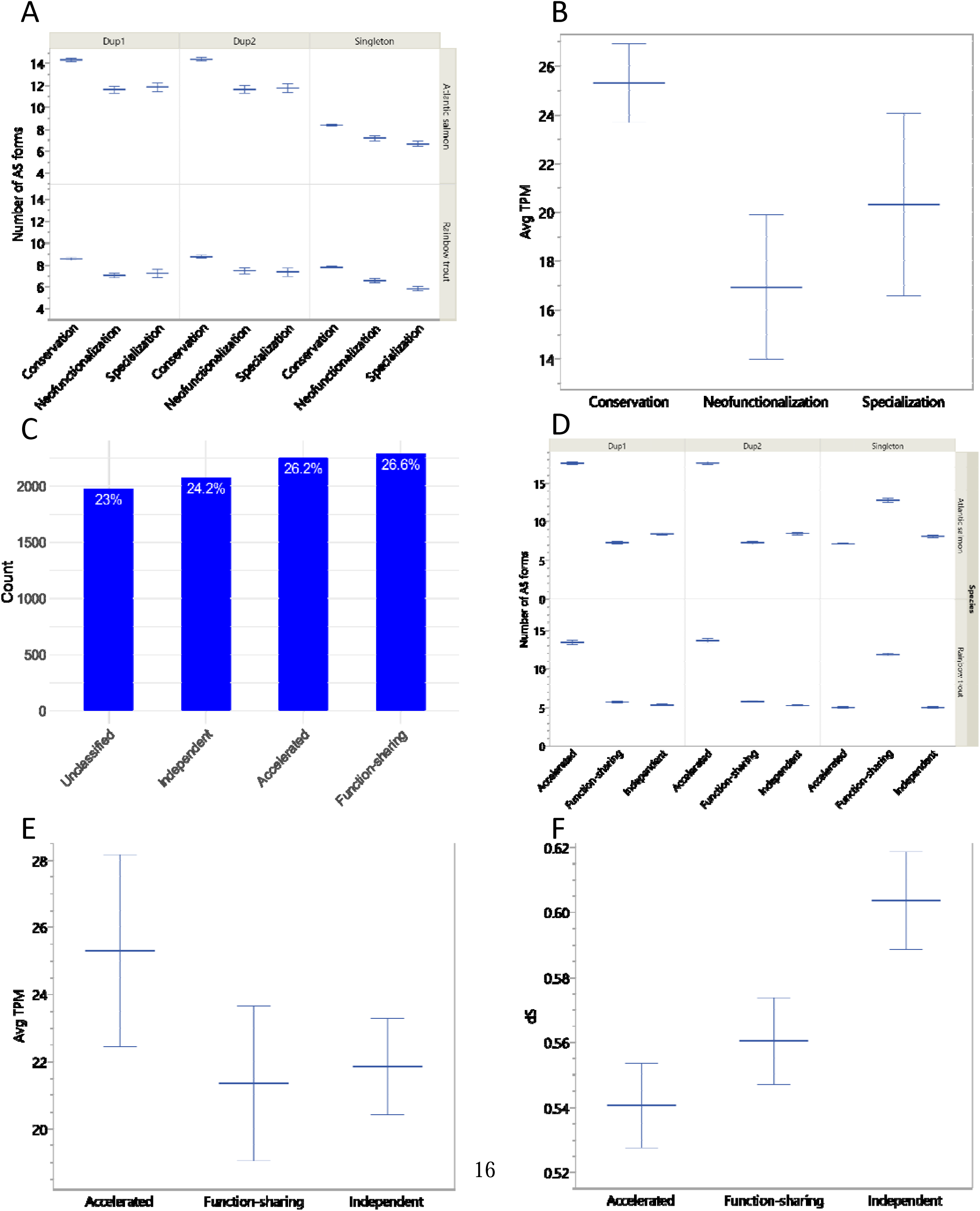
(A) Number of AS forms in conserved, neofunctionalized, and specialized gene duplicates (D1 & D2) and their single-copy orthologs. Genes with low levels of ancestral splicing tend to diverge in expression following WGD event. (B) Conserved genes display significantly higher levels of expression than neofunctionalized and specialized gene duplicates. (C) Count and percentage of gene pairs that belongs to each AS evolutionary model. The data supported a combined scenario of the accelerated AS, independent, and function-sharing models (D) Number of AS forms in gene triplets (D1-D2-singleton) of each AS evolutionary model. For the accelerated model, gene duplicates have more splicing forms than their singleton orthologs in the Northern Pike. Gene pairs supporting the function-sharing model exhibit reduced AS forms, which are about half as abundant as the ancestral splicing forms. In the independent model, singletons and gene duplicates have a similar number of AS forms. (E) Genes that undergo accelerated splicing display significantly higher levels of expression than those supporting the function-sharing and independent models. (F) The average dS value of genes that experience accelerated splicing is significantly lower than that of independent genes, suggesting that ohnologues tend to lose AS forms with the evolutionary time.

#### Evolutionary models of alternative splicing after WGD

To identify the primary evolutionary model of AS, we clustered the ohnologue pairs based on the splicing level of the unduplicated sister taxon (i.e., Northern pike). First, ohnologue pairs with more AS forms than the ancestral singletons would support the evolutionary accelerated AS model. Second, ohnologue pairs with fewer AS forms than the ancestral singletons would support the evolutionary function-sharing model. Third, ohnologue pairs with similar AS forms to the ancestral singletons would support the evolutionary independent model. We found 26.2% of the tested gene duplicates in rainbow trout showing accelerated splicing compared to the ancestral singletons (Figure 5C & D). The sum of the accelerated AS forms of the rainbow trout ohnologues exhibited a moderate correlation (R = 0.61) with the ancestral state in Northern pike. In the meantime, a comparable number of the gene duplicates (26.6%) revealed fewer splice variants than their ancestral singletons supporting the function-sharing model (Figure 5C & D). The sum of the reduced AS forms of the rainbow trout ohnologues is highly correlated with the ancestral state in Northern pike (R = 0.88). In Atlantic salmon, a higher percentage of gene duplicates (∼46.9%) supported the accelerated model, whereas ∼8.7% supported the function-sharing model. Notably, a significant proportion of the Arlee trout and salmon gene duplicates supported the independent model with AS forms similar to those of the Northern Pike singletons. We observed a highly significant difference in the expression levels between genes that undergo accelerated splicing and genes that undergo either function-sharing (Wilcoxon test, P < 4.3E-41) or independent splicing (Wilcoxon test, P < 5.5E-22) (Figure 5E), suggesting that splicing dynamics may be tightly linked to gene function.

Data from both rainbow trout and Atlantic salmon highlight a significant role for the independent model in the evolution of splicing forms following WGD. This finding challenges the conclusions drawn by a previous study[11], which had dismissed the independent model’s role in the evolutionary process of teleost’s AS. Notably, the lower average dS values (synonymous substitution rates) in genes undergoing splicing acceleration compared to independent genes (Figure 5F) suggest that splicing acceleration may be associated with stronger selective constraints. This finding supports the current study’s conclusion that ohnologues gradually lose AS forms over evolutionary time. Additionally, it introduces the independent model as the primary long-term evolutionary model of AS following WGD events.

#### Divergence of epigenetic regulatory elements between different modes of splicing and gene expression

Epigenetic signatures are mainly enriched in the promoters and enhancers. It has become increasingly known that these signatures also exist in the gene bodies, implying a potential relationship between epigenetic regulation and pre-mRNA splicing[23]. Splicing is a co-transcriptional process in which chromatin histone modifications affect the final splicing product. It has been proposed that chromatin modifications recruit chromatin-binding proteins to modulate the splicing factors binding to pre-mRNAs[24].

Therefore, we hypothesized that gene types with significant differences in the proportion of AS forms would also have distinct epigenetic signatures. To test this hypothesis, ChIP-Seq data generated from six tissues (brain, liver, muscle, intestine, kidney, and spleen) were used to identify histone marks in the promoter (2Kb upstream of TSS) and gene bodies (Figure 6A-D). We focused on four histone marks: H3K4me3, which correlates with promoters of active genes[25–28]; H3K27me3, which is associated with promoters of inactive genes[29, 30], H3K27ac which is a mark of active regulatory elements and promoters[30]; and H3K4me1, which is correlated with enhancers[30]. The WGD duplicates exonic and intronic regions revealed higher proportions of all histone tags than other gene types (single-copy genes and dispersed duplicates). H3K27ac and H3K4me1 were the most abundant histone marks in the WGD duplicates (Figure 7A&B). The exonic and intronic regions of ohnologues undergoing accelerated splicing revealed a higher abundance of H3K27ac, H3K4me1, and H3K4me3 than ohnologues supporting function-sharing and independent models (Wilcoxon test, P < 7.08E-41) (Figure 7C). Similar patterns of the H3K27ac, H3K4me1, and H3K4me3 were also observed in 2kb upstream of the TSS (Wilcoxon test, P = 0) (Figure 7C). Conversely, the presence of H3K27me3 was exclusively found to be lower upstream of the TSS of genes undergoing accelerated splicing (Wilcoxon test, P < 5.18E-08) (Figure 7C). The H3K27me3 profiles of gene pairs with accelerated splicing were less correlated than those of genes that undergo sharing of AS forms and independent splicing (Wilcoxon test, P < 0.0003) (Figure 7D), suggesting a potential regulatory role of H3K27me3 in the splicing process of these genes. Overall, the study highlights a distinct correlation between epigenetic signatures and AS in WGD duplicates. Specifically, ohnologues undergoing accelerated splicing exhibit higher levels of active histone marks, such as H3K27ac, H3K4me1, and H3K4me3, across both exonic and intronic regions, as well as upstream promoter regions, compared to those supporting function-sharing and independent models. Conversely, the repressive mark H3K27me3 is significantly reduced in these regions for genes with accelerated splicing, suggesting its potential role in modulating splicing outcomes. These findings emphasize the complex interplay between chromatin modifications and splicing regulation, indicating that distinct epigenetic landscapes may contribute to differential splicing patterns in WGD duplicates. The study also supports the hypothesis that gene types with varied AS forms display unique epigenetic signatures, advancing our understanding of how chromatin dynamics influence splicing evolution post-WGD.

**Figure 6:**
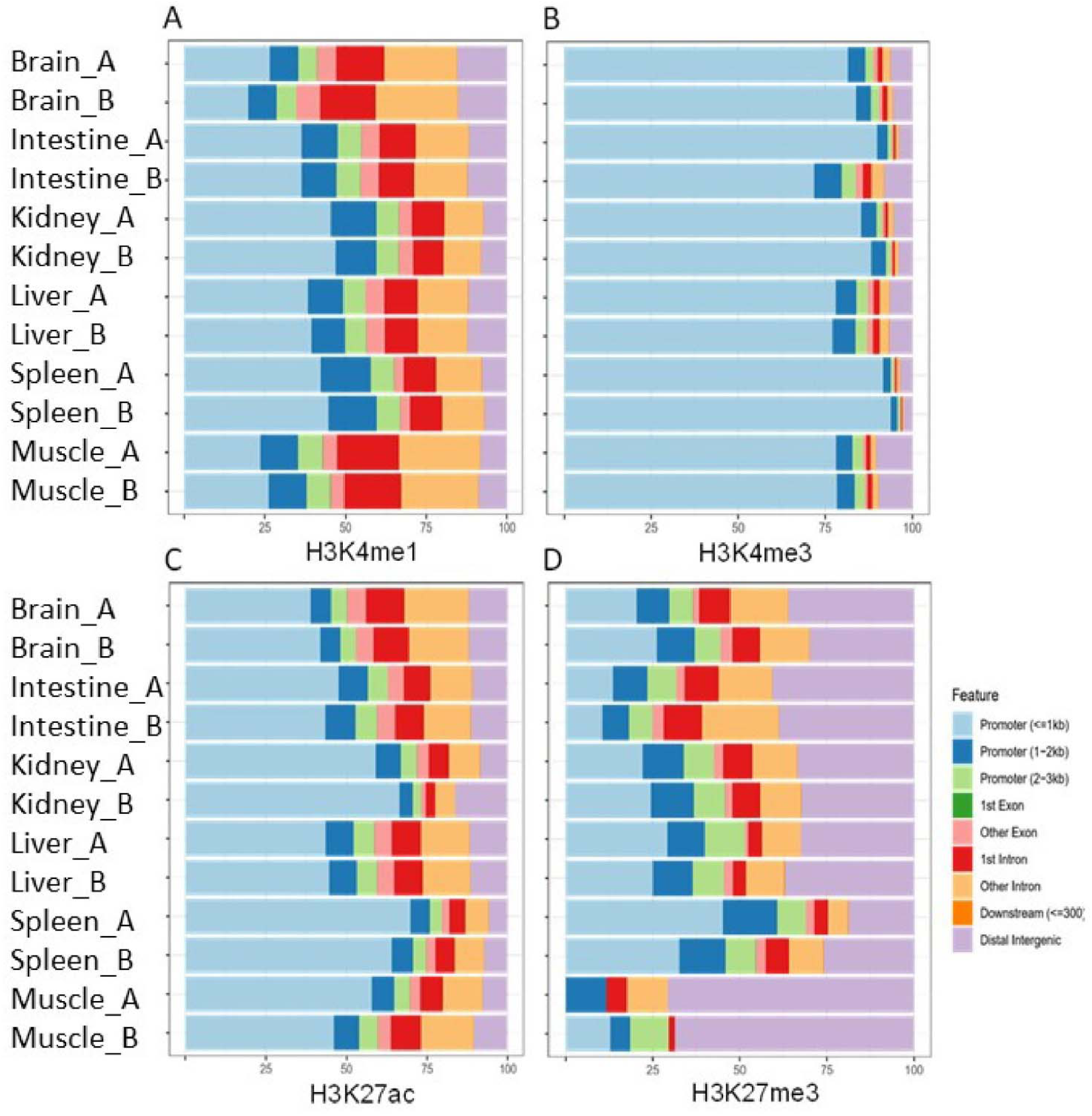
(A-D) Bar plots display the percentage of histone peaks (H3K4me1, H3K4me3, H3K27ac, and H3K27me3) that overlap with various genomic features, including the TSS, exonic, intronic and intergenic regions.

**Figure 7:**
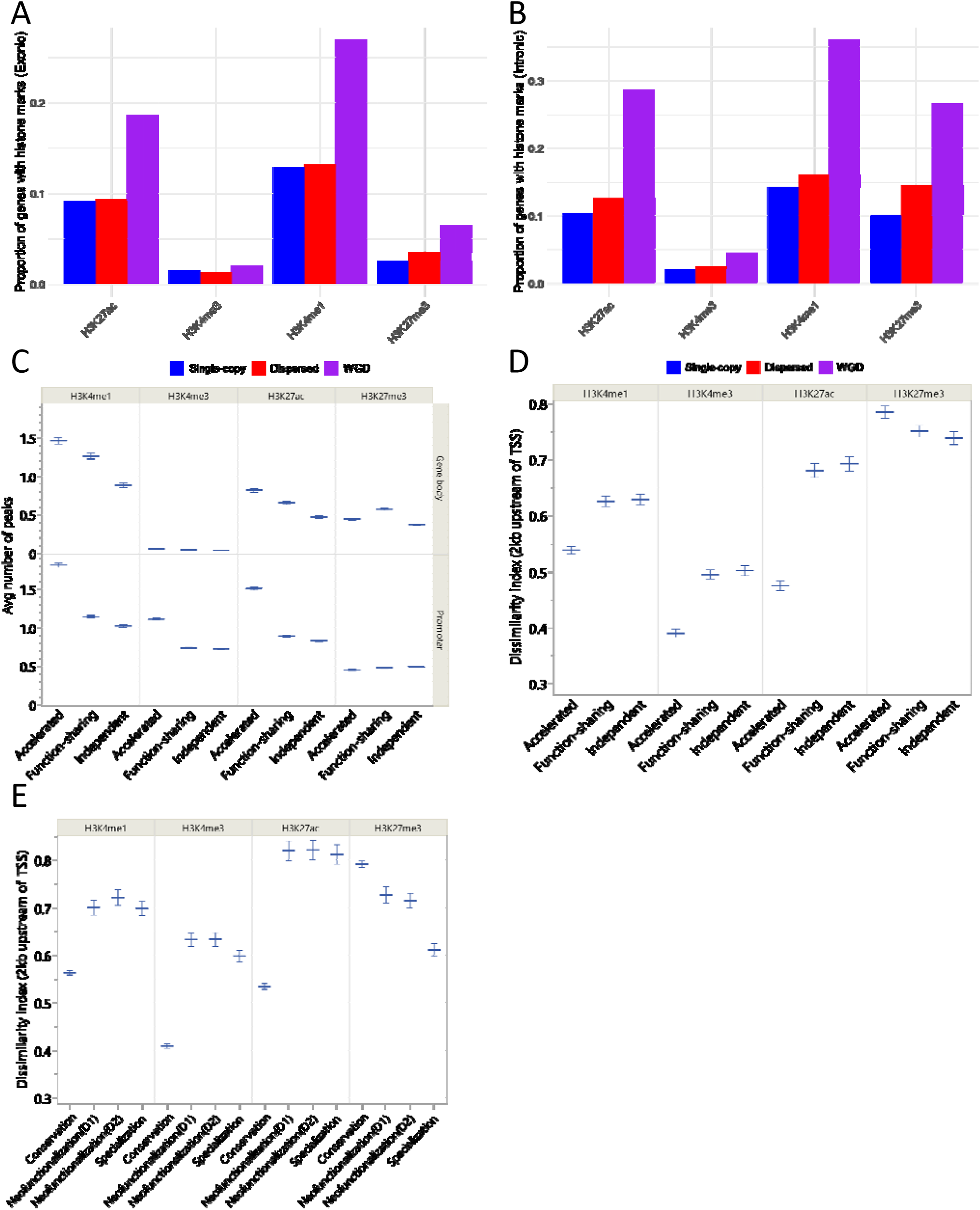
(A & B) ChIP-Seq data from six tissues identified a significantly higher representation of histone marks in the gene body of WGD duplicates than other gene types. (C) Gene pairs exhibiting accelerated splicing have more active enhancer-associated histone marks (H3K4me1 and H3K27ac) 2Kb upstream of the TSS and in the gene body, which may help explain their increased splicing efficiency. (D & E) The histone modification divergence of each category’s duplicate gene pair was assessed using the dissimilarity index (1-r), where r is the Pearson correlation coefficient of the histone modification profiles for the duplicated gene pair. Gene pairs with conserved expression or accelerated splicing exhibit the least divergence of the active enhancer– and promoter-associated histone marks profiles upstream of TSS.

We further analyzed the divergence in histone modification profiles between conserved, neofunctionalized, and specialized ohnologues (Figure 7E). Divergence was assessed using a dissimilarity index based on the histone modification profiles in the promoter regions (see Materials and Methods). Our results show that, compared to neofunctionalized genes, conserved gene paralogues display significantly less divergence in their histone modifications (Wilcoxon test, P < 2.05E-139). This lower divergence in conserved paralogues aligns with their stable gene expression patterns, indicating a close relationship between histone modification conservation and gene expression. In contrast, the higher divergence observed in neofunctionalized genes reflects their evolving regulatory mechanisms, supporting the idea that epigenetic divergence plays a key role in driving functional innovation in these genes.

## DISCUSSION

It is well known that, among other factors, protein diversity increases from AS, gene duplication, and the accumulation of mutations. Previous studies have focused on mammalian species and a few other model organisms, including non-salmonid teleost fish, to elucidate the relationship between gene duplications and AS[8–11]. Since salmonids experienced a recent WGD, their genomes contain many ohnologues (WGD duplicates), which offers an excellent opportunity to study the evolution of gene expression and AS divergence early after duplication. Studies performed on salmonids focused on investigating gene expression to gain insights into the consequence of WGD events in adaptive genome evolution and mechanisms that maintain the retention of duplicate genes[2, 5, 6, 21].

Divergence in expression levels of duplicate genes plays a crucial role in the evolution of novel/complex phenotypes[4, 5, 18, 21]. Paralogs retained from WGD (ohnologues) are typically associated with the organism’s development and have been reported to be prone to dominant deleterious mutations and involved in genetic diseases[31]. However, the evolutionary mechanisms driving the functional evolution of duplicate gene copies remain uncertain in many species[18]. Despite multiple chromosomal rearrangements since the salmonids’ 4^th^ WGD, the rainbow trout genome has remained relatively conserved[21]. Consistently, 24,336 (62.6%) duplicated genes were inferred from collinear blocks of the current rainbow trout genome assembly. Such a unique advantage of salmonid genomes can give insight into the early evolutionary fate of duplicated genes after WGD. Despite 100 Mya of evolution since the Ss4R event, more than 70% of the ohnologues were maintained through conservation, suggesting conservation is a crucial step in facilitating the initial survival of duplicate genes. Remarkable preservation of the expression of the two ancestral genome copies was previously detected in rainbow trout[21]. Compared to our study, Lien et al.[5] found that only 42% of the Ss4R duplicates exhibited conserved co-expression with their orthologs in Northern pike. Our study also discovered distinct epigenetic signatures associated with the conserved genes. Except for H3K27me3, conserved gene paralogues displayed lower divergence in their histone modification profiles than neofunctionalized genes. H3K27me3 has been reported to significantly disrupt the transcriptional robustness of highly conserved genes in fungi[32]. Among the histone marks studied, H3K4me3 of the conserved gene pairs displayed the highest correlation, surpassing H3K4me1, H3K4me3, and H3K27me3, which aligns with the conservation of gene expression. Such conservation for about 100 Mya disagrees with previous studies reporting rapid divergence of the gene expression levels of duplicate genes[12–15].

Discrepancy in ranking individual genes within duplicate pairs, based on correlation to their orthologs, has raised questions regarding the conclusions drawn from previous studies investigating the preservation of gene duplicates[4–7]. Therefore, we sought to revisit the debate over the relative importance of sub– and neo-functionalization in the genome’s evolution after WGD events. We used the Euclidian distances method[17, 18], not Pearson’s correlation coefficient, which revealed robustness to gene ranking. Consistent with previous reports from Atlantic salmon[5, 6], we found that subfunctionalization of expression is rare and that a considerable proportion of the duplicate genes are neofunctionalized. Neofunctionalized genes exhibit higher dS values, indicating that they may evolve novel functions due to relaxed selective constraints. This observation suggests that neofunctionalization may serve as the long-term mechanism for retaining duplicates. In agreement with this observation, we noticed a lower proportion of conserved genes accompanied by a higher proportion of neofunctionalized genes in three teleost fish species compared to salmonids. This trend was particularly evident in medaka, which possesses a relatively small genome of approximately 700 Mb. These findings disagree with Braasch et al.[4], who proposed that partitioning ancestral gene functions rather than the evolution of new gene functions is the potential mechanism of duplicate gene preservation. Additionally, our study sheds light on a distinct group of gene pairs that have evolved differently from their ancestral orthologs. Lien et al.[5] have proposed the possibility of subfunctionalization within this particular cluster. Here, we propose that these genes constitute a specialized cluster in which the gene pairs are distinct from each other and from the ancestral ortholog. Notably, the neofunctionalized and specialized gene pairs’ H3K27ac profile exhibited the most significant dissimilarity compared to the H3K4me3 profile of the conserved genes, aligning with the observed divergence in gene expression. These results indicate a substantial role for the enhancer in the functional divergence between ohnologues following WGD. A recent study on Atlantic salmon by Verta et al. (2021) reported a correlation between transcriptional divergence in duplicated genes resulting from WGD and variations in the number of nearby regulatory elements. This finding suggests that enhancers primarily drive the functional divergence between ohnologues following WGD[33].

This study shows a more complex relationship between AS and gene duplication than previously thought[8, 9]. We demonstrate that type of gene duplication, divergence time, and ancestral splicing confound the complex relationship. We then determined the evolutionary models of AS after WGD in the Swanson line of rainbow trout and validated the main conclusion on the Arlee line of rainbow trout and Atlantic salmon. Our results showed more than one evolutionary model of AS after WGD, making a combination of multiple models more likely. However, our analyses of the rainbow trout data revealed that the accelerated model is likely the initial evolutionary model of AS after the WGD event. It was previously reported that this model proposes a relaxed selection pressure for each paralogous gene, leading to increased AS per gene (reviewed in[34]). Su et al. (2006) supported the function-sharing model immediately after gene duplication[9]. We present evidence that ohnologues gradually lose splice variants over time under relaxed purifying selection pressure. This result contradicts the generic claim of immediate loss of AS forms in recently duplicated genes[9]. The finding also aligns with the observation that gene pairs belonging to the function-sharing and independent models have higher average dS values than the accelerated model. Further, our results indicate that conserved genes with the lowest average of dS values and the highest level of ancestral splicing tend to possess more AS forms than specialized and neofunctionalized genes. This suggests a substantial role for AS in balancing the gene dosage to ameliorate immediate fitness costs posed by genome duplication. Genes with low levels of ancestral splicing tend to diverge in expression following the WGD event, suggesting a crucial role for these genes in triggering the evolution of novel adaptive phenotypes.

Furthermore, this work elucidates a significant role played by the independent model in the evolution of splicing forms following WGD. In contrast to previous reports[9, 11], our findings reveal a high prevalence of independent splicing instances in salmonids. It was previously suggested that functional divergence among human duplicate genes at the genome level could potentially decrease the number of AS forms. This finding aligns more closely with the predictions of the function-sharing model rather than the independent model[9]. Additionally, Wang and Guo have ruled out the independent model in zebrafish, medaka, and stickleback[11]. The current study demonstrates that genes that undergo independent splicing have a significantly higher mean of dS values than genes splicing through acceleration. This observation suggests that the independent model is likely the driving force for the long-term evolution of AS following WGD. Of note, genes undergoing function-sharing and independent splicing revealed similar epigenetic signatures to neofunctionalized and specialized genes, which suggests a substantial role for enhancers in driving the AS evolution following WGD.

Overall, dosage balance, which may not directly involve the evolution of new functions in the short term, may pave the way for eventual neofunctionalization and, thus, retention of duplicate genes. Besides, this study not only presents compelling evidence for the initial evolution of AS through acceleration but also highlights the crucial role of the independent model in driving its long-term evolution. In conclusion, retaining duplicates may trigger the evolution of new adaptive functions, allowing the occupation of new ecological niches.

## METHODS

### Homology search and gene classification

Protein sequences and genomic positions of a newly assembled and annotated rainbow trout genome (Accession GCA_025558465.1) was used for this study. Additionally, protein sequences and genomic positions for other fish species were mainly retrieved from the Ensembl genome browser (Atlantic salmon “GCA_905237065.2”, Northern pike “GCA_004634155.1”, Japanese medaka “GCA_002234675.1”, three-spined stickleback “BROADS1”, Atlantic cod “GCA_902167405.1”, zebrafish “GCA_000002035.4”, Mexican tetra “GCA_000372685.2”, chinook salmon “GCA_002872995.1”, coho salmon “GCA_002021735.2”, and spotted gar “GCA_000242695.1”). Also, the RefSeq annotation of Atlantic salmon “GCF_905237065.1” was included in the analysis. The transcript with the longest CDS was used for genes with more than one transcript. Protein-coding genes from each species were compared against themselves and those of other species using BLASTp (i.e., All-vs.-All local BLASTp) to search for homology. The best five non-self-hits for each protein sequence in each target genome(s) with an E-value threshold < 10^−5^ were reported for each protein sequence.

The MCScanX software package[35] was used to classify genes of each species into five types based on their copy number and genomic distribution; single-copy, dispersed duplicates, tandem duplicates, proximal duplicates, and WGD/segmental duplicates. The BLASTp output and annotation file were used as the input files to execute the duplicate gene classifier, a core program of MCScanX. Classes of gene duplication were determined according to the following: all genes were initially labeled as singletons and assigned ranks according to their order on chromosomes. Genes with BLASTp hits to other genes were re-labeled as dispersed duplicates. Gene pairs were re-labeled as proximal duplicates when they had a difference of gene rank < 20 (configurable) or tandem duplicates if the difference of gene rank = 1. Finally, the MCScanX was executed, and anchor genes in collinear blocks were re-labeled as segmental/WGD duplicates. When a gene appeared in multiple hits, it was assigned to a unique class according to the following order of priority: WGD/segmental>tandem>proximal>dispersed.

### Gene families

For the Swanson genome, we only kept the longest isoform expressed from each gene locus. All sequences were used to build a database for BLAST. All-by-all BLASTp was performed to find the top hit for each sequence in the database. The clustering program, MCL[36], was used to group the most similar sequences into gene families. Single-copy genes identified by both MCScanX and MCL were used for the downstream analyses.

### Evolutionary models of alternative splicing

Under the null hypothesis of no difference in the number of AS forms, we applied the statistical test suggested by Pocock[37] to compare the number of AS forms of two groups (*p*-value < 0.05). For the accelerated model, a significant difference was expected between the number of AS forms in Northern Pike and the sum of the number of AS forms in trout’s duplicate gene pairs. Whereas a statistically nonsignificant difference was expected in the case of the function-sharing model. Also, when the number of AS forms of the two groups is expected to be similar, we required statistical nonsignificant differences.

### Calculation of non-synonymous and synonymous substitution rates

For each gene class, protein sequences of each gene pair were aligned using MUSCLE[38]. The amino acid alignments were converted into nucleotide alignments using pal2nal[39], with –nogap argument. The alignment files were then converted to AXT format to calculate the Ka (dN) and Ks (dS) substitution rates, and the Ka/Ks ratios using the KaKs Calculator 3.0[40].

### ChIP-Seq analyses

The raw sequencing data was subjected to quality control using FastQC (version v0.11.8)[41]. Adapter sequences were removed from the reads using trim galore (version 0.6.4) (https://github.com/FelixKrueger/TrimGalore). Trim galore utilizes Cutadapt internally to trim adapter sequences and performs additional quality filtering steps to improve data quality. Trimmed reads were aligned to the rainbow trout reference genome using Bowtie2 (version 2.3.5.1)[42] with the “--end-to-end” and “--very-sensitive” parameters. These parameters enabled end-to-end alignment and increased sensitivity, reducing false alignments. The aligned reads were stored in BAM format for subsequent analysis. MACS3 (Model-based Analysis of ChIP-Seq 3) (version 3.0.0a6)[43] was used for peak calling. The aligned reads in BAM format were utilized as input, and specific parameters were adjusted for each histone modification. For H3K27me3, which represents broad peaks, we set the q-value threshold to 0.05. For the rest of the histone modifications, which represent narrow peaks, we set the q-value threshold to 0.01. These thresholds were used to identify significant peaks, representing regions of enrichment associated with the respective histone modifications. ChIPseeker (version 1.36)[44] was employed for peak annotation. The identified peaks were annotated by assigning them to genomic features, such as genes, promoters, enhancers, and other relevant regions.

### Divergence of Histone modifications

We calculated the log2-transformed fold enrichment ratio. Subsequently, we transformed these ratios into z scores using the formula Z_X_ = (χ − μ)/δ. Here, χ represents the ratio value for a given gene, μ denotes the mean ratio of all genes, and δ signifies the standard deviation of this ratio across all genes. To evaluate the divergence of histone modification patterns between duplicate gene pairs, we employed the dissimilarity index, which is calculated as “1− r”. In this equation, r represents the Pearson correlation coefficient of the histone modification profiles for the duplicated gene pair. We compared the mean values of the dissimilarity index in each gene category and the significance was determined using the Wilcoxon rank-sum test.

### RNA-seq and quantification of gene expression

The raw RNA-seq reads for rainbow trout (Acc# SRP108798 and ERP003742), Atlantic salmon (Acc# SRP011583), Northern pike (Acc# SRP040114 and SRP045141), zebrafish (Acc# SRP044781), medaka (Acc# SRP044784), and spotted gar (Acc# SRP044782) were downloaded from NCBI SRA. The raw reads were trimmed using CLC Genomics Workbench (version 22.0). High-quality reads were mapped to the reference genome sequence of the corresponding species using the HISAT2 aligner[45]. The abundance levels of each gene were retrieved from the BAM files using the TPMCalculator (https://github.com/ncbi/TPMCalculator). The unique reads were used to measure the gene expression levels in different tissues.

### Identification of gene Triplets and mechanisms of duplicate gene preservation

Salmonid WGD duplicates, retrieved from the output file containing collinear blocks identified by the MCScanX[35], were blasted against singletons from the Northern pike. When both members of the duplicate gene pair hit the same singleton (*E*-value < 10^−5^), the gene triplet was considered for the downstream analysis. Additionally, WGD duplicates from non-salmonid fish (pike, zebrafish, and medaka) were blasted against singletons from the spotted gar to identify gene triplets.

We restricted the analyses to triplets for which each gene copy is expressed in at least one tissue. The expression profile of the singletons in the Northern pike or spotted gar was used as a proxy for the expression prior to duplication. All absolute expression levels were converted to relative expression levels (proportions of contributions to total expression). The relative expression values were used as gene expression profiles for comparison. To control for the sex, RNA-Seq datasets from male Swanson, Atlantic salmon, and Northern pike were used to investigate the fate of duplicate gene pairs in salmonids. Whereas datasets from female spotted gar and non-salmonid teleost fish were used to investigate the long-term preservation of duplicate genes.

We classified the evolutionary processes/mechanisms preserving pairs of duplicate genes by applying the phylogenetic method developed by Assis and Bachtrog[17] to expression profiles of D1, D2, and singletons. We calculated Euclidian distances between expression profiles of D1 and ancestral copies (*E _D1,A_*), between expression profiles of D2 and ancestral copies (*E _D2,_*_A_), and between the combined D1– D2 expression profile and that of the ancestral copy (*E _D1+D2,A_*). The baseline gene divergence level was then established by calculating Euclidian distances between expression profiles of singletons in sister species (*E* _S1,S2_). Several cutoff values were explored to define expression divergence. We selected the semi-interquartile range from the median due to its robustness to outliers. Finally, each pair of duplicates was classified as conserved, neofunctionalized, subfunctionalized, or specialized according to previously established rules[17]. We expect *E* _D1,A_[≤[*E* _S1,S2_ and *E* _D2,A_[≤[*E* _S1,S2_ when duplicates are conserved, *E* _D1,A_[>[*E* _S1,S2_ and *E* _D2,A_[≤[*E* _S1,S2_ when the D1 is neofunctionalized, *E* _D1,A_[≤[*E* _S1,S2_ and *E* _D2,A_[>[*E* _S1,S2_ when the D2 is neofunctionalized, *E* _D1,A_[>[*E* _S1,S2_, *E* _D2,A_[>[*E* _S1,S2_, and *E* _D1+D2,A_[≤[*E* _S1,S2_ when duplicates are subfunctionalized, and *E* _D1,A_[>[*E* _S1,S2_, *E* _D2,A_[>[*E* _S1,S2_, and *E* _D1+D2,A_[>[*E* _S1,S2_ when duplicates are specialized.

### Ethics

A laboratory strain of double-haploid rainbow trout was sampled according to the Standard Operating Procedures: Care and Use of Research Animals. Protocol Number 114, USDA, ARS National Center for Cool and Cold Water Aquaculture, 11861 Leetown Road, Kearneysville, WV 25430, USA. The Institutional Animal Care and Use Committee reviewed and approved this protocol on November 17, 2016.

## Data access

All raw and processed sequencing data generated in this study have been submitted to the NCBI Gene Expression Omnibus (GEO; https://www.ncbi.nlm.nih.gov/geo/) under accession number GSE245212. https://www.ncbi.nlm.nih.gov/geo/query/acc.cgi?acc=GSE245212,

Ensembl and RefSeq accessions of all genome assemblies used in the current study were reported in the methods section. Similarly, we provided the NCBI SRA accession codes of the raw RNA-Seq reads that were downloaded from the NCBI Sequence Read Archive (SRA) to calculate unique TPM.

Computational pipelines: https://github.com/arealia/Evolution-of-Rainbow-Trout-Genome

## Acknowledgements

Liqi An, Ying Wang, Xuechen Bai, and Ye Bi, from Huaijun Zhou laboratory are acknowledged for their help with the generating the ChIP-seq work for the rainbow trout genome annotation, data were reused from original data of a complete study that will be published somewhere else.

## Proofreading

Grammarly (2025) was used for text improving and proofreading[46].

