## Supplementary_Tables_and_Figs_4-25 for "Splicing Retention and Enhancer Divergence Govern the Evolutionary Fate of Ohnologues Following Whole-Genome Duplication in Rainbow Trout": Supplementary_Tables_and_Figs_4-25.pdf

**Table S1:** Classification of evolutionary processes maintaining rainbow trout WGD duplicates, resulting from application of various cutoffs to Euclidian distances. Trout's gene expression data were retrieved from **Salem et al. (2015)[1]**. **Random** order was given to the duplicate gene copies (i.e., labels duplicate 1 "D1" and duplicate 2 "D2" were randomly designated to the two genes).

| E_div | Value | Conservation | Neofunctionalization(D1) | Neofunctionalization(D2) | Subfunctionalization | Specialization |
|---|---|---|---|---|---|---|
| MeanSD | 0.798 | 7226 | 445 | 469 | 9 | 447 |
| mean2SD | 1.084 | 8294 | 115 | 121 | 2 | 64 |
| medSIQR | 0.636 | 6134 | 734 | 730 | 29 | 969 |
| MedIQR | 0.827 | 7367 | 405 | 432 | 14 | 378 |
| quant75 | 0.682 | 6484 | 662 | 635 | 20 | 795 |

**SUMMARY:** Herein, we classified the evolutionary processes essential for the preservation of rainbow trout WGD duplicates. We applied various cutoffs to Euclidian distances and randomly assigned labels (D1 and D2) to the gene pairs. While all the cutoff values showed that duplicate genes are mainly maintained by conservation (at least 71.4%), a considerable fraction of gene pairs were maintained by neofunctionalization and specialization (3.5-28.3%). Our findings revealed a rare occurrence of subfunctionalization following WGD.

**Table S2:** Classification of evolutionary processes maintaining rainbow trout WGD duplicates, resulting from application of various cutoffs to Euclidian distances. Trout's gene expression data were retrieved from **Salem et al. (2015)[1]**. Duplicate gene pairs were **ranked** as "most conserved" (i.e., the gene copy with the highest expression correlation with the ortholog in the outgroup; D1) and "most diverged" (i.e., the gene copy with the lowest expression correlation with the ortholog in the outgroup; D2).

| E_div | Value | Conservation | Neofunctionalization(D1) | Neofunctionalization(D2) | Subfunctionalization | Specialization |
|---|---|---|---|---|---|---|
| MeanSD | 0.798 | 7226 | 87 | 827 | 9 | 447 |
| mean2SD | 1.084 | 8294 | 48 | 188 | 2 | 64 |
| medSIQR | 0.636 | 6134 | 133 | 1331 | 29 | 969 |
| MedIQR | 0.827 | 7367 | 95 | 742 | 14 | 378 |
| quant75 | 0.682 | 6484 | 120 | 1177 | 20 | 795 |

**SUMMARY:** Ranking the individual genes within the duplicate pairs[2] resulted in two extreme conclusions of a previous study[3] investigating the mechanism of gene duplicate preservation. To test whether the ranking would affect our conclusion, we ranked the gene pairs as "most conserved" (i.e., the gene copy with the highest expression correlation with the ortholog in the outgroup; D1) and "most diverged" (i.e., the gene copy with the lowest expression correlation with the ortholog in the outgroup; D2) as in [2]. Interestingly, our approach was robust to the ranking process and yielded the same results obtained from random ranking (Table S3).

**Table S3:** Classification of evolutionary processes maintaining rainbow trout WGD duplicates, resulting from application of various cutoffs to Euclidian distances. Trout's gene expression data were retrieved from **Berthelot et al. (2014)[4]**. **Random** order was given to the duplicate gene copies (i.e., labels duplicate 1 "D1" and duplicate 2 "D2" were randomly designated to the two genes).

| E_div | Value | Conservation | Neofunctionalization(D1) | Neofunctionalization(D2) | Subfunctionalization | Specialization |
|---|---|---|---|---|---|---|
| MeanSD | 0.771 | 7205 | 473 | 415 | 7 | 452 |
| mean2SD | 1.069 | 8221 | 127 | 124 | 3 | 77 |
| medSIQR | 0.586 | 6086 | 770 | 708 | 22 | 966 |
| MedIQR | 0.782 | 7263 | 457 | 400 | 8 | 424 |
| quant75 | 0.645 | 6538 | 651 | 596 | 16 | 751 |

**SUMMARY:** Duplicate genes may show tissue-specific, time-point-specific, or simultaneous divergent expression[5]. Therefore, we sought to replicate our conclusion (Table S3) using RNA-seq data from fish used to assemble the trout[4] (BioProject PRJEB4450) genome. Consistent with the results above in Table S3, more than 70% of duplicate genes were preserved by conservation whereas 28.6% of gene pairs were maintained by neofunctionalization and specialization.

**Table S4:** Classification of evolutionary processes maintaining **Atlantic salmon** WGD duplicates, resulting from application of various cutoffs to Euclidian distances. Salmon's gene expression data were retrieved from RNA-Seq datasets downloaded from the NCBI SRA (BioProject PRJNA72713). **Random** order was given to the duplicate gene copies (i.e., labels duplicate 1 "D1" and duplicate 2 "D2" were randomly designated to the two genes).

| E_div | Value | Conservation | Neofunctionalization(D1) | Neofunctionalization(D2) | Subfunctionalization | Specialization |
|---|---|---|---|---|---|---|
| MeanSD | 0.782 | 9410 | 489 | 527 | 7 | 485 |
| mean2SD | 1.082 | 10512 | 135 | 174 | 6 | 91 |
| medSIQR | 0.606 | 8239 | 826 | 830 | 28 | 995 |
| MedIQR | 0.803 | 9515 | 451 | 496 | 7 | 449 |
| quant75 | 0.657 | 8651 | 712 | 729 | 22 | 804 |

**SUMMARY:** We also sought to replicate our conclusion (Tables S3 & S5) in a closely-related species. For this purpose, RNA-seq data from Atlantic salmon[6] (BioProject PRJNA72713) were used to retrieve the gene expression values. Consistent with the results above in Tables S3 & S5, more than 70% of duplicate genes were preserved by conservation whereas 24.3% of gene pairs were maintained by neofunctionalization and specialization.

**Table S5:** Classification of evolutionary processes maintaining rainbow trout WGD duplicates, resulting from application of various cutoffs to Euclidian distances. Trout's gene expression data were retrieved from **Salem et al. (2015)[1]**. Only **housekeeping genes** that displayed RPKM values $\geq 1$ in each tissue of both the Northern pike and rainbow trout were used.

| E_div | Value | Conservation | Neofunctionalization(D1) | Neofunctionalization(D2) | Subfunctionalization | Specialization |
|---|---|---|---|---|---|---|
| MeanSD | 0.439 | 1041 | 14 | 98 | 0 | 59 |
| mean2SD | 0.562 | 1168 | 2 | 26 | 0 | 16 |
| medSIQR | 0.361 | 836 | 13 | 188 | 5 | 170 |
| MedIQR | 0.424 | 1012 | 17 | 115 | 0 | 68 |
| quant75 | 0.364 | 856 | 9 | 179 | 5 | 163 |

**SUMMARY:** To further validate our results and eliminate any potential bias caused by the developmental, physiological, and immunological conditions of the studied fish, we reanalyzed the gene expression data were retrieved from **Salem et al. (2015)[1]** using housekeeping genes that displayed RPKM values $\geq 1$[1] in each tissue of both the Northern pike and rainbow trout (BioProject PRJNA389609). Consistent with the whole dataset, a similar pattern was also achieved when only housekeeping genes were used (69% conservation, 16.6% neofunctionalization, 14.0% specialization, and 0.4% subfunctionalization).

**Table S6:** Classification of evolutionary processes for long-term preservation of WGD duplicates, resulting from application of medSIQR cutoff to Euclidian distances. Gene expression levels were computed from RNA-Seq datasets generated from Northern pike, zebrafish, and medaka.

| | medSIQR | Conservation | Neofunctionalization(D1) | Neofunctionalization(D2) | Subfunctionalization | Specialization |
|---|---|---|---|---|---|---|
| Trout-Salem et al.[1] | 0.636 | 6134 | 734 | 730 | 29 | 969 |
| Trout-Berthelot et al.[4] | 0.586 | 6086 | 770 | 708 | 22 | 966 |
| Salmon-Lien et al.[6] | 0.606 | 8239 | 826 | 830 | 28 | 995 |
| Northern pike | 0.552 | 1100 | 215 | 223 | 15 | 214 |
| Zebrafish | 0.572 | 418 | 70 | 92 | 3 | 77 |
| Medaka | 0.578 | 443 | 117 | 114 | 5 | 91 |

**SUMMARY:** Compared to salmonid species, we observed reduced proportions of conserved genes accompanied by increased proportions of neofunctionalized and specialized genes in non-salmonid species, including Northern pike, zebrafish, and medaka.

### SUPPLEMENTARY FIGURES

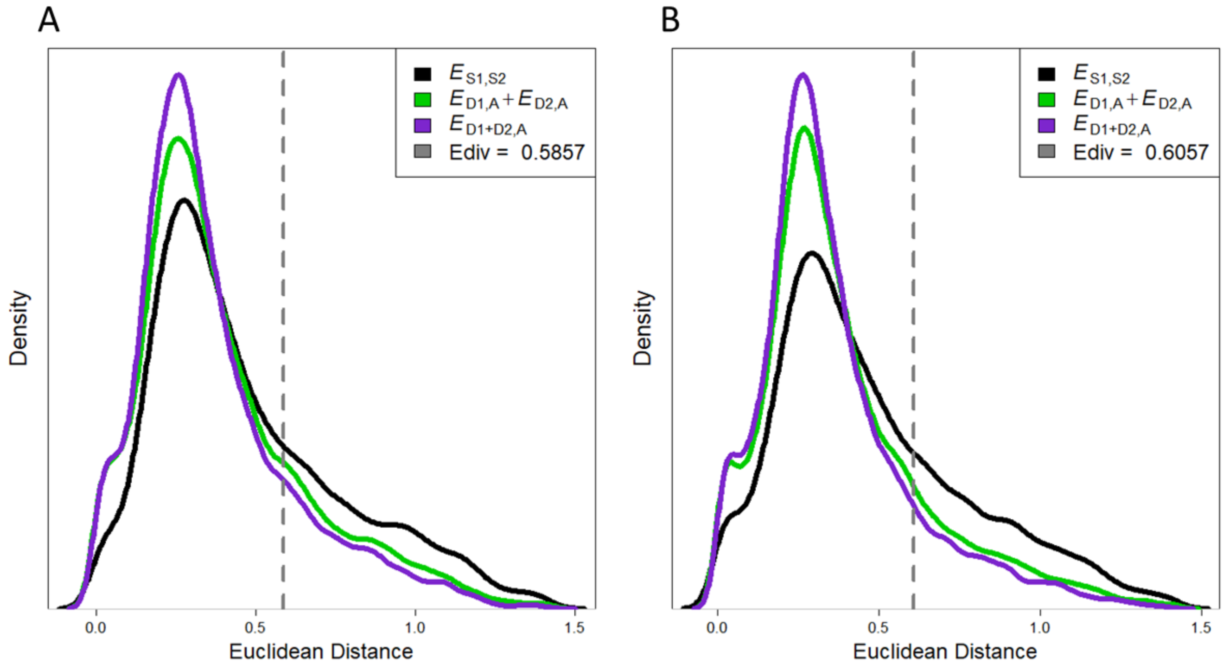


**Figure S1:** Distributions of all computed Euclidian distances and the position of $E_{div}$. Gene expression data from Berthelot et al. (2014)[4] for WGD duplicate genes of rainbow trout were used for the analysis. Similarly, gene expression data from Lien et al. (2016)[6] (B) for WGD duplicate genes of Atlantic salmon were used for the analysis. $E_{D1,A}$ + $E_{D2,A}$ is a single distribution for $E_{D1,A}$ and $E_{D2,A}$. The plots in panels A and B suggest that most of the salmonid gene duplicates are retained by conservation.

**SUMMARY:** Herein, we investigated the evolutionary processes essential for the preservation of rainbow trout and Atlantic salmon WGD duplicates using data produced by Berthelot et al. (2014)[4] (Panel A) and Lien et al. (2016)[6] (Panel B). We applied the medSIQR cutoff to Euclidian distances and randomly assigned labels (D1 and D2) to the gene pairs. The plots in panels A & B suggest that most of gene duplicates are retained by conservation. Results from this analysis align with those obtained using data generated by Salem et al. (2015) and presented in the main manuscript.

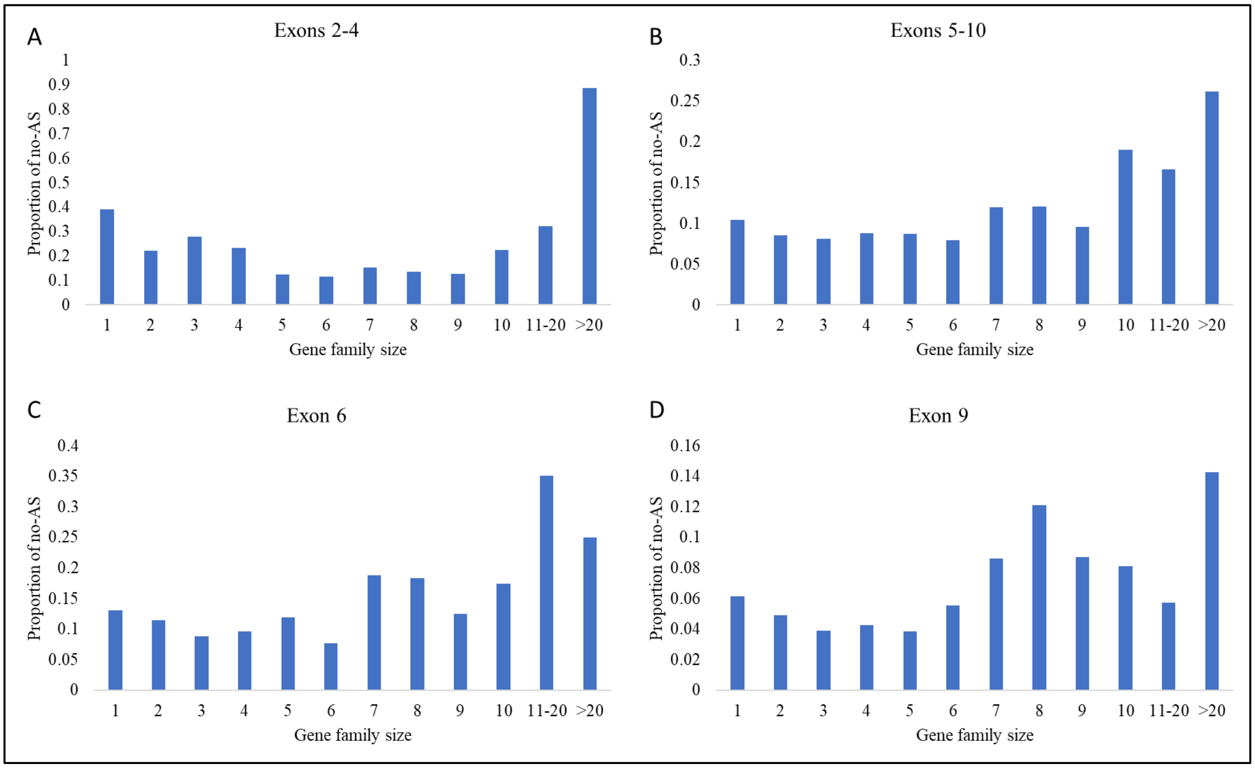


**Figure S2:** Relationship between the proportion of no-AS trout genes and the size of the gene family. Four exon interval bins were generated, and the single-copy genes did not always show the lowest proportion of no-AS, particularly compared to gene family sizes 2 to 6, in all the tested bins.

**SUMMARY:** Exon duplication was reported as a mechanism for the evolution of AS[7]. In order to ensure that our findings are not due to the relationship between the exon counts and AS, we re-evaluated the relationship between gene family size and AS using specific intervals of exon counts. For intervals of less than 10 exons, we noticed that singletons did not show the highest proportion of AS in all the tested scenarios. Also, using the median (n = 6) and average (n = 9) exon numbers for singletons, the latter tended to show a lower proportion of AS forms.

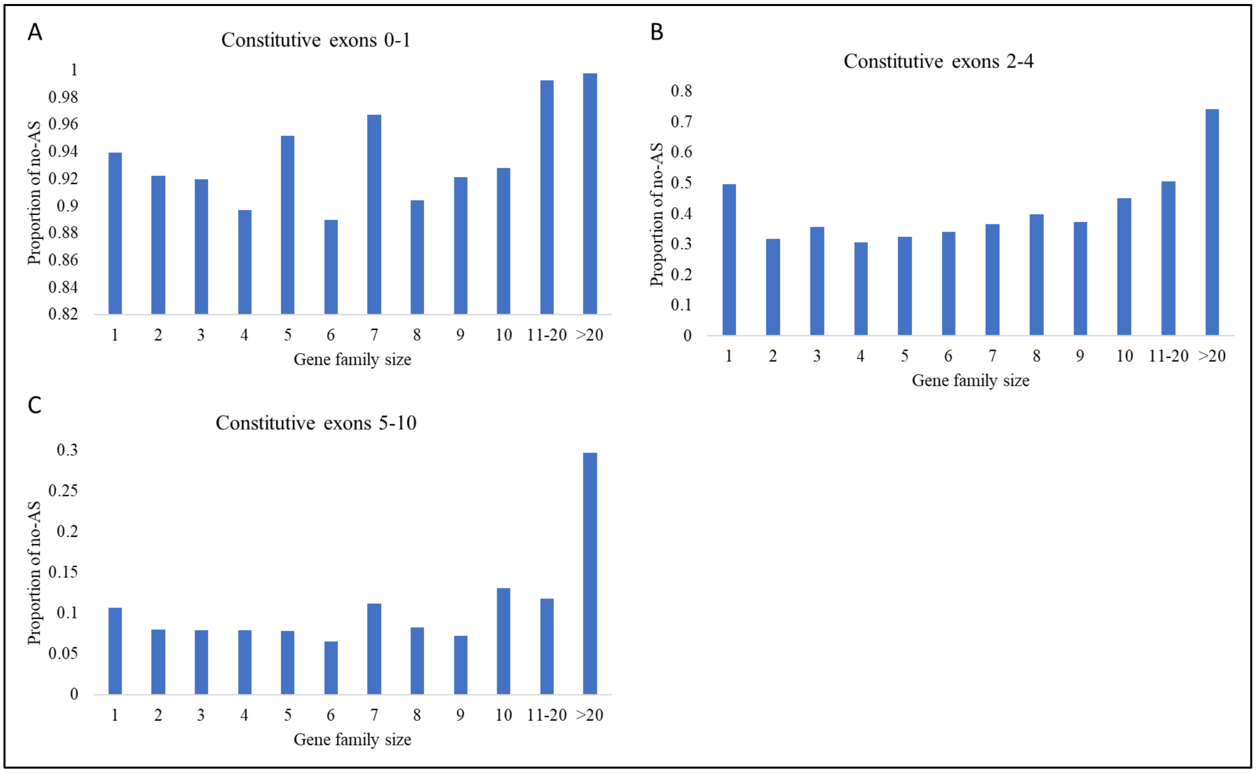


**Figure S3:** Relationship between the proportion of no-AS trout genes and the size of the gene family. Three bins of constitutive exons were generated, and the single-copy genes did not always show the lowest proportion of no-AS in all the tested bins.

**SUMMARY:** A previous study reported that genes with fewer constitutive exons are faster in acquiring splice forms[8]. Therefore, we sought to control for the number of constitutive exons. For intervals of less than 10 constitutive exons, we also noticed that singletons did not show the highest proportion of AS in all the tested scenarios.

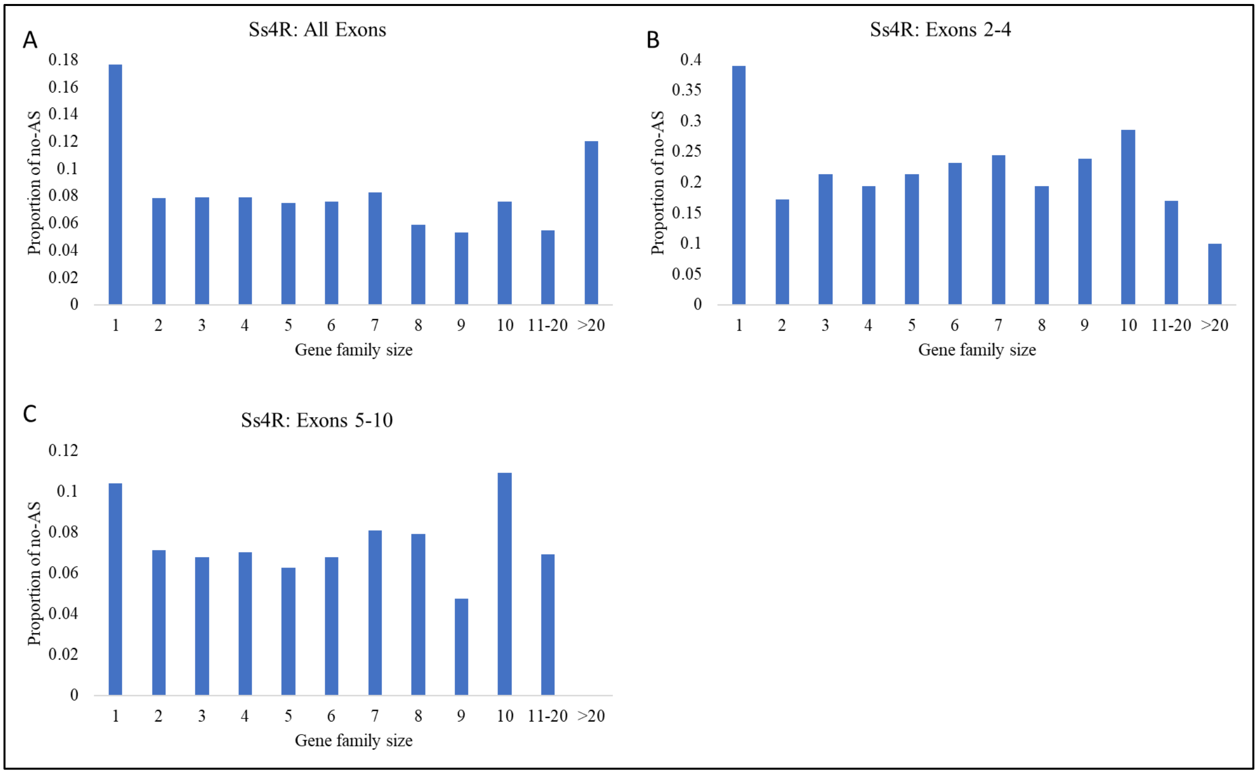


**Figure S4:** (A) The relationship between the proportion of no-AS forms of Ss4R genes and gene family size. The proportion of no-AS forms of all duplicated gene families was less than that of the single-copy gene family. (B & C) Two exon interval bins were generated, and the single-copy genes exhibited the lowest proportion of no-AS compared to most of the duplicated gene family sizes in both tested bins.

**SUMMARY:** Ss4R ohnologues revealed lower proportions of no-AS than singletons in all gene family sizes. Re-evaluating the relationship between the proportion of no-AS forms of Ss4R genes and gene family size in two bins of exon intervals (2-4 and 5-10) revealed also lower proportions of no-AS in the Ss4R ohnologues. Our results ensure that the observed relationship between the proportion of no-AS forms of Ss4R genes and gene family size are not due to the relationship between the exon counts and AS.

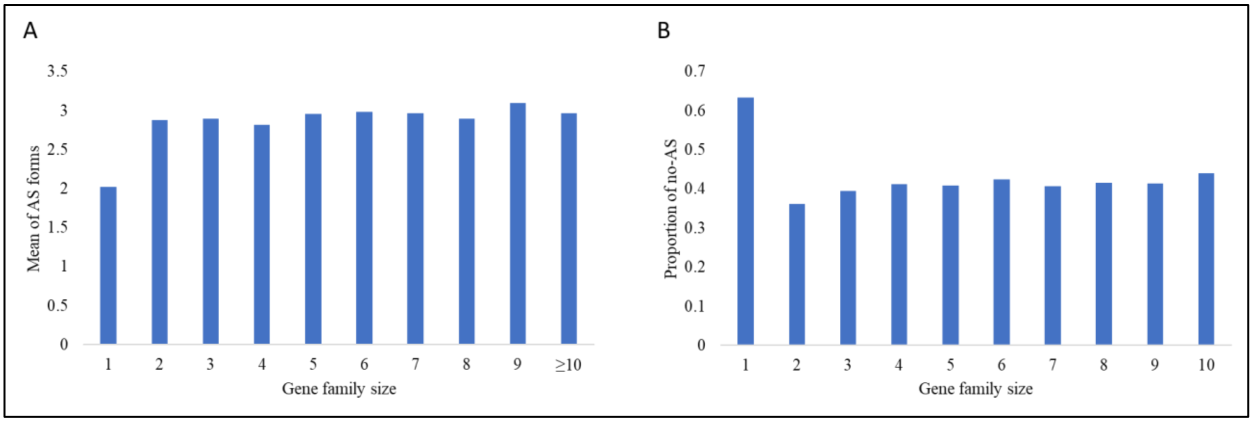


**Figure S5:** (A) Using Iso-Seq data, the mean of AS forms of all duplicated gene families was higher than that of the single-copy gene family. (B) There was a higher proportion of no-AS in singletons compared to all duplicated gene families (i.e gene family size ≥ 2).

**SUMMARY:** In this study, we sought to re-evaluate the relationship between the proportion of AS forms and gene family size using PacBio long-reads with splice junctions supported by short reads from our previous study [9] to avoid potential false positive splice junctions introduced by the gene prediction tool "Augustus" or assembled RNA-Seq transcripts. Using only long reads, we observed fewer AS forms in singletons than in small gene family sizes. These results prove that the observed relationship between the proportion of AS forms and gene family size is not due to a specific methodological choice.

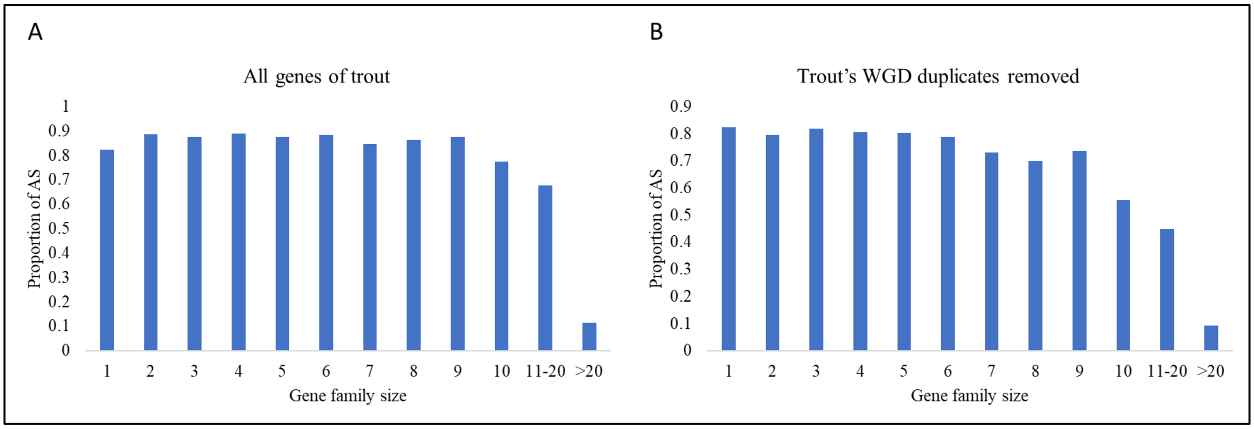


**Figure S6:** (A) A parabola curve was observed between the proportion of AS forms and gene family size. The proportion of AS forms in duplicated gene families, except gene family size ≥ 10, was higher than in the single-copy gene family. While there is a higher proportion of alternatively spliced genes in larger gene families, a remarkable increase in the proportion of the genes with no-AS compared to singletons was observed only in gene family sizes ≥10 (B) The relationship between the proportion of AS forms and gene family size when WGD duplicates are absent. A remarkable decline in the fraction of AS forms was noticed while the gene family size increases.

**SUMMARY:** Using the whole set of genes to investigate the relationship between the proportion of AS forms and gene family size, a parabola curve was observed with lower levels of AS in single-copy genes and genes with family sizes above 10, as opposed to those between 2-10. By removing WGD duplicates from the whole pool of genes used for the analysis, a remarkable decline in the fraction of AS forms was noticed while the gene family size increased. Gene families 2 to 10 exhibited a strong negative correlation with the proportion of AS forms (R = -82.1, *P* = 5.13E-05). Single-copy genes had the highest proportion of AS forms compared to other gene family bins.

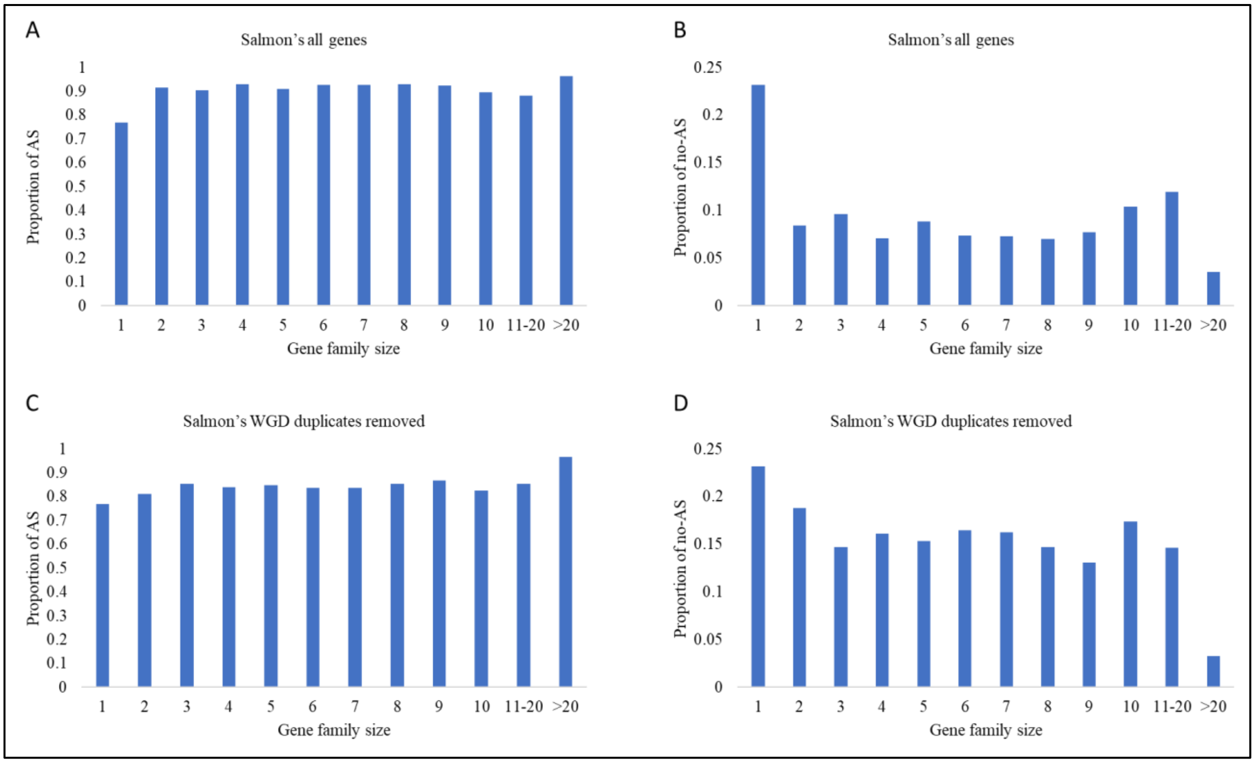


**Figure S7:** (A) The relationship between the proportion of AS forms and gene family size in Atlantic salmon. The proportion of AS forms of all duplicated gene families was higher than that of the single-copy gene family. (B) A higher proportion of no-AS forms was observed in singletons than in all duplicated gene family sizes. (C & D) Removing WGD duplicates of Atlantic salmon led to dramatic decrease in proportion of AS and increase in the proportion of no-AS forms.

**SUMMARY:** In Atlantic salmon, the proportion of no-AS in singletons is significantly higher than that of all large-size duplicate gene family sizes (≥ 2). By removing the WGD duplicates, we found that the proportion of no-AS forms significantly increased in gene family sizes 2 up to 20 (Wilcoxon Signed-Rank Test $P$ = 0.004), but did not exceed that of the singletons.

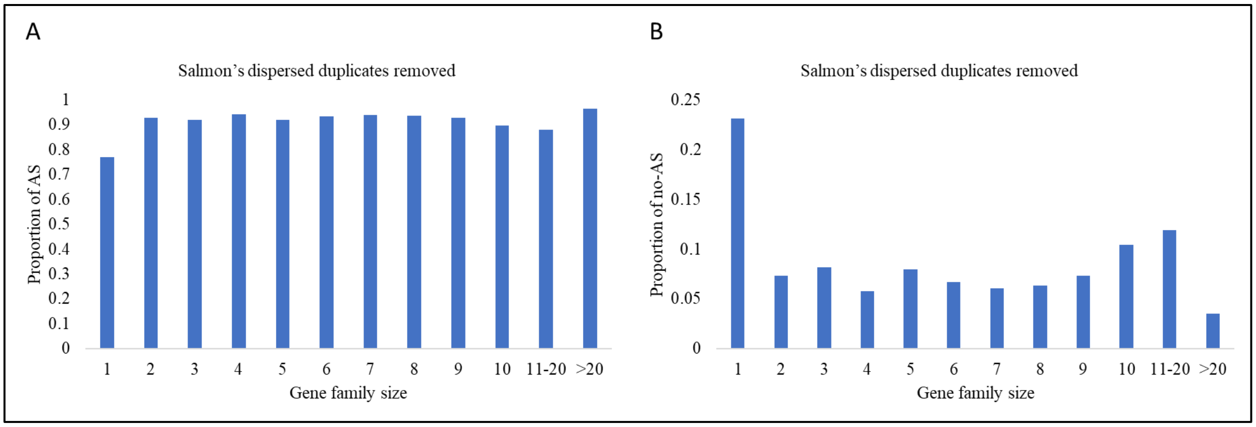


**Figure S8:** (A) The relationship between the proportion of AS forms and gene family size in Atlantic salmon after removing all dispersed genes. Removing dispersed genes significantly increased the proportion of AS forms of duplicated gene families. (B) A significant reduction in the proportion of no-AS forms of duplicated gene families following removal of dispersed genes.

**SUMMARY:** Contrary to WGD, removing dispersed duplicates of Atlantic salmon from the analysis resulted in a significant reduction in the proportion of no-AS forms (Wilcoxon Signed-Rank Test $P < 0.05$).

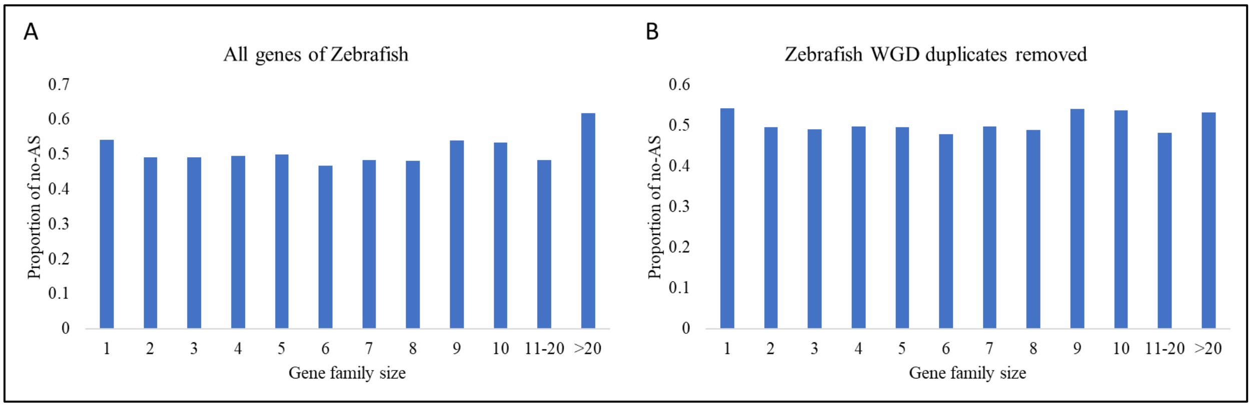


**Figure S9:** (A) The relationship between the proportion of no-AS forms and gene family size in zebrafish. (B) Removing WGD duplicates from zebrafish gene families did not significantly impact the proportion of no-AS forms.

**SUMMARY:** In zebrafish, the proportion of no-AS in singletons is higher than that of large-size duplicate gene family sizes (2-20). The proportion of no-AS forms was 0.54 and 0.49 in singletons and gene family size 2, respectively. By removing the WGD duplicates, we found that the proportion of no-AS forms did not significantly increased in gene family sizes 2 up to 20 (Wilcoxon Signed-Rank Test $P > 0.05$). For instance, we observed the same proportion of no-AS forms (0.49) in gene family 2.

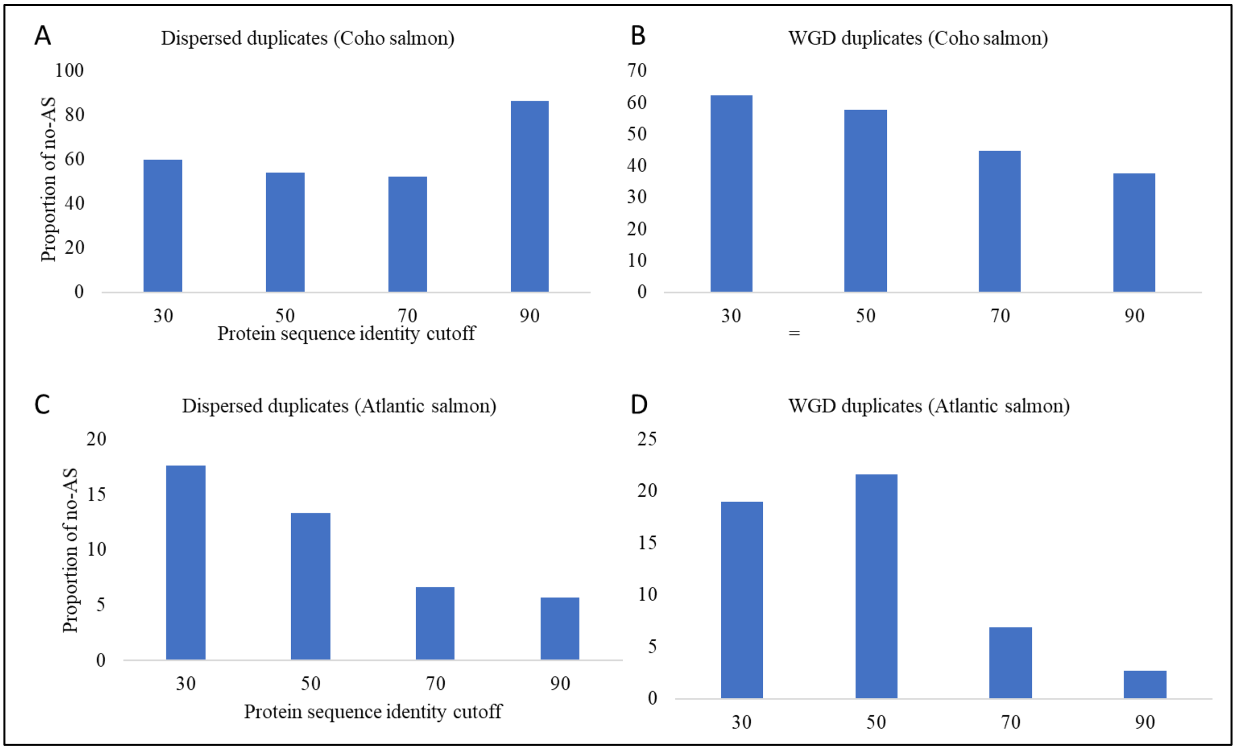


**Figure S10:** In coho salmon, recent dispersed duplicates are less likely to be alternatively spliced (A), whereas WGD duplicates tend to lose AS forms as they evolve over time (B). In Atlantic salmon (Ensemble annotation), both dispersed duplicates (C) and WGD duplicates tend to lose AS forms under relaxed purifying selection pressure (D). Each bar represents the fraction of genes not subjected to AS. Group 30 consisted of genes showing a sequence identity ranging from 30% to 50% ($30 \leq I < 50$) at an E-value less than 1E-5. Similarly, the groups 50, 70, and 90 had sequence identity within the ranges of $50 \leq I < 70$, $70 \leq I < 90$, and $I \geq 90$, respectively.

**SUMMARY:** We used the amino acid identity percentage ($I$), at an E-value less than1E-5, from the self-genome BLASTp output to group the duplicate genes of coho salmon and Atlantic salmon into four bins ($30 \leq I < 50$, $50 \leq I < 70$, $70 \leq I < 90$, and $I \geq 90$). We then calculated the proportion of no-AS genes in each bin. The amino acid identity ($I$) was used as a proxy for divergence time. For WGD duplicates of coho salmon, we found that as $I$ increases, the proportion of no-AS genes decreases, which means loss of AS forms does not happen shortly after the gene duplication. Conversely, we found that that as $I$ increases, the proportion of no-AS dispersed duplicate genes increases, which means the loss of AS isoforms in dispersed duplicates happens shortly after the gene duplication. However, in Atlantic salmon, the dispersed duplicates, annotated by the Ensembl, unexpectedly revealed a similar pattern to the WGD duplicates. Contrary to the Ensembl-annotated dispersed duplicates, the RefSeq annotation showed that the proportion of dispersed genes with no-AS forms in the Atlantic salmon increased as $I$ increases (Figure 5). Our analysis reveals that the loss of AS forms is contingent upon the divergence time and gene duplication type (WGD or dispersed duplicates).
